# HLA Alleles Imprint Distinct Biases in the Usage Preferences of TCR Vβ segments

**DOI:** 10.64898/2026.02.19.706729

**Authors:** Leonardo V. Castorina, Matthew T. Noakes, Lorenzo Pisani, Julia Greissl, Harlan Robins, Haiyin Chen-Harris, H. Jabran Zahid

## Abstract

T cells must co-recognize peptide and HLA, yet the extent to which this specificity is shaped by germline-encoded TCR–HLA contacts versus selection during thymic development has remained difficult to quantify. Leveraging population-scale TCR*β* repertoires linked to donor HLA genotypes, we construct allele-specific V*β* usage profiles and normalize them to repertoire-wide baselines to derive interpretable HLA-TCR V*β* preference vectors. We demonstrate that different HLA alleles imprint distinct biases in the usage frequency of TCR V*β* gene segments among the public TCRs that engage those alleles; consistent with germline-encoded TCR–HLA contact preferences, certain V*β* genes are over-represented for particular HLA alleles. Similarities in HLA amino-acid sequence predict similarities in both their V*β* preferences and peptide-binding motifs; a residue-level analysis disentangles HLA positions primarily associated with TCR engagement from those associated with peptide motifs. The TCR-associated HLA positions localize to canonical TCR-facing helices, whereas peptide-associated HLA positions track binding pockets, revealing distinct molecular routes by which HLA polymorphism shapes the TCR and peptide sides of recognition. CMV exposure stratifications confirmed these patterns are not explained by a few dominant infections. Together, these data support that germline-encoded constraints set the landscape of TCR–HLA compatibility, while thymic and peptide-driven forces tune the realized repertoire. These HLA-specific V*β* biases are a biological prior that should provide a baseline for better understanding of TCR-pHLA specificity as a whole and should be accounted for in future evaluation of any TCR-pHLA specificity prediction methods.

## 1 Introduction

Interactions between T cell receptors (TCRs) and peptide-presenting Human Leukocyte Antigens (HLAs) are central to adaptive immunity, enabling the immune system to distinguish self from non-self and mount targeted responses to pathogens, cancer and other immunological threats (Murphy and Weaver, 2017a). A deeper understanding of this interaction could transform our ability to predict immune responses, design vaccines, personalize immunotherapies and uncover the pathogenesis of autoimmune disorders. Achieving this will likely require a precise understanding of how TCRs engage with peptide–HLA (pHLA) complexes at the molecular level. Despite substantial progress (Davis and Bjorkman, 1988; Rudolph et al., 2006; Rossjohn et al., 2015; Murphy and Weaver, 2017b), key aspects of this molecular interaction remain incompletely understood.

Understanding TCR–pHLA interactions is complicated by their extraordinary complexity. HLAs are among the most polymorphic human genes, with tens of thousands of alleles identified across the six classical class I (HLA-A, -B, -C) and II loci (HLA-DR, -DQ and -DP) (Klein and Sato, 2000). Structurally, the HLA peptide-binding groove is defined by two helices: in Class I the helices are named *α*1 and *α*2 and are part of the same polypeptide chain, while for Class 2 the groove is formed by two distinct chains, *α*1 and *β*1. In the canonical binding mode, the TCR*β* interacts with around the *α*1 helix in Class I and the *β*1 helix in Class II (See Figure5 A-D) Christopher Garcia et al. (2009); Cohn et al. (2019)).

Polymorphisms are concentrated in the peptide-binding groove, influencing both peptide presentation and TCR binding (Tiercy, 2002; Nakamura et al., 2019; Karnaukhov et al., 2022). Moreover, TCR binding is degenerate: a single TCR can recognize many pHLAs and many TCRs can recognize the same pHLA (Davis and Bjorkman, 1988; Sewell, 2012; Murphy and Weaver, 2017b). While many predictive models use sequence-based, structural or molecular dynamics approaches (Springer et al., 2021; Grazioli et al., 2023; Bradley, 2023; Jensen and Nielsen, 2024), they often fail to generalize (Drost et al., 2025), limited by interaction complexity and sparsity of high-quality data (Grazioli et al., 2022; Castorina et al., 2025). In the absence of massive training sets, uncovering mechanistic principles can provide inductive biases that improve model generalizability. This strategy mirrors the success of AlphaFold, which leveraged Multiple Sequence Alignments to identify co-evolutionary constraints and guide structural prediction (Jumper et al., 2021). Similarly, statistical frameworks like ours may uncover residue-level features that constrain TCR–pHLA binding and inform the next generation of predictive models. This perspective is especially relevant given ongoing efforts to determine whether TCR specificity arises from germline-encoded features such as CDR1 and CDR2 loops that may have evolved HLA specificity or is shaped entirely by thymic selection acting on random repertoires (La Gruta et al., 2018). Clarifying these mechanisms may guide the development of more general and interpretable models.

Recent advances in sequencing technologies have enabled statistical approaches to examine how HLA polymorphism shapes TCR gene usage. Sharon et al. (2016) proposed that if germline-encoded contacts influence TCR–HLA interactions, then TCR V*β*s should show biased usage depending on HLA type. Supporting this, their trans-eQTL analysis found that polymorphisms near the HLA binding groove correlate with V*β* expression, linking HLA polymorphism to TCR repertoire composition. However, their use of bulk RNA sequencing averaged over the entire T cell compartment, potentially dilutes the HLA-specific signals. To improve resolution, we leverage a previously developed method that uses large-scale repertoire sequencing to associate public TCR*β*s (i.e., those shared across individuals) with specific HLAs. This approach, introduced and validated by Zahid et al. (2025a), leverages TCR co-occurrence across thousands of subjects with known HLA genotypes to identify TCR*β*s statistically associated with specific HLAs. These associations generalize to diverse, unseen populations, yielding TCR*β* sets with strong evidence of true immunological specificity.

Using HLA-associated TCR*β* sets, we quantify V*β* gene usage for each HLA and examine how these patterns correlate with HLA polymorphism, while statistically accounting for potential mediation by shared peptide motifs. We identify HLA residues associated with V*β* usage, peptide repertoire or both. To compare patterns across HLA types, we harmonize sequence and structural data through sequence alignments and 3D structure mapping. Although our analysis is limited to the TCR*β* chain, it reveals population-level trends consistent with germline influences and highlights candidate HLA residues that may shape TCR–HLA specificity through direct structural mechanisms. The scale of this analysis enables resolution of TCR–HLA associations not previously accessible, offering biologically grounded priors to inform predictive models.

## 2 Materials and Methods

### 2.1 HLAdb

Using direct HLA genotyping and 4,144 TCR*β* repertoires, Zahid et al. (2025a) built sensitive and specific HLA imputation models from TCR*β*s associated with hundreds of common HLAs. We impute 145 of the most common HLAs for 30,000 subjects in the T-Detect COVID cohort (Zahid et al., 2025b). These imputed HLAs were used as case-control labels and TCR*β*s were statistically associated to HLAs using via a one-sided Fisher’s Exact Test following Zahid et al. (2025a). By imputing HLAs on a set of repertoires that are nearly an order of magnitude larger in sample size than the original data set used to build the models, we are able to leverage the genotyped labels available for a smaller set of repertoires and the statistical power of sequence discovery offered by the larger cohort.

We restricted analysis to HLAs with more than 5,000 associated public TCR*β*s. In total, 2,827,857 public TCR V*β* sequences from 41 unique public TCR V*β* genes associated with 91 HLA alleles were included in this analysis (see Supplementary Table S1). Using the set of TCR*β*s associated with each HLA, we generate the TCR V*β* usage pattern as the distribution of V*β* gene preferences for that particular HLA. V*β* genes are not uniformly represented in the TCR*β* repertoire, thus we normalize V*β* gene usage to the background frequency measured from all TCR*β*s (public and private) summed across repertoires and this is represented as a vector (see Supplementary Figure S2 and Supplementary Table S1).

### 2.2 TCR V*β* Normalization

We start with a matrix whose rows are HLAs and columns are TCR V*β* s, containing co-occurrence counts. We also have background counts for each TCR V*β* . First, we remove ambiguous or undefined V*β* genes. We then normalize TCR*β* V-gene counts by dividing by the corresponding background counts to correct for any underlying baseline TCR V*β* frequency, for each TCR V*β* gene, the total number of times it was observed across 30,000 repertoires was used for background normalization. Any ambiguous or undefined V*β* genes were removed.

After background adjustment, we convert counts into frequencies by dividing each each *N*_*ij*_ by the total for HLA *i*:

- For each HLA *i*, compute 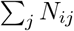 .
- Define 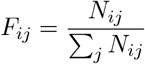

This ensures 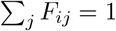 for each HLA *i*.

We call the frequency vector *F*_*i*_ for HLA *i* the “V*β* gene preference”.

### 2.3 Pairwise Distance Analysis of HLA

The TCR-pHLA interaction involves the TCR, the HLA and a peptide. We characterize HLA differences with three pairwise distance matrices:

- **HLA polymorphism**: BLOSUM62 distance between HLA sequences (See 2.3.1).
- **V***β* **gene preference change**: cosine distance between V*β* preference vectors (See 2.3.2).
- **Peptide motif change**: Jensen–Shannon (JS) divergence between Position Weight Matrix (PWM) (See 2.3.3).

We then correlate these matrices to identify HLA positions whose variation is associated with TCR*β* usage or peptide motifs, disentangling TCR–HLA from pHLA effects.

We analyse both **within** HLA loci and **between** loci. Within-locus analyses aim to identify binding-groove positions most significant for the peptide or the TCR*β*.

Between-locus analyses aim to identify sequence positions that distinguish loci and correlate with the TCR*β* or the peptide motifs.

#### 2.3.1 HLA Polymorphism

We use the HLA sequences dataset from the TCR Dock pipeline Bradley (2023), restricted to the TCR-binding groove to ensure comparable sequence lengths and also focuses the analysis around areas of major polymorphism.

HLA sequences are aligned using Clustal *ω* (Sievers et al., 2011). Pairwise differences are computed using the BLOSUM62 substitution matrix (Henikoff and Henikoff, 1992) whose log-odds quantify amino acids substitution likelihoods. We calculate both per-position and overall differences by summing scores across positions..

For position *k* in aligned sequences *S*_*i*_ and *S*_*j*_, we calculate a per-position distance, *D*_*seq*_[*k*], using the exponential of the BLOSUM score:

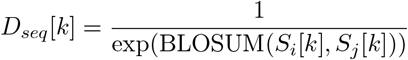

The overall sequence divergence is calculated by summing these per-position distances:

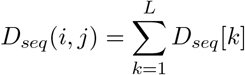

To summarize:

- *S*_*i*_, *S*_*j*_: HLA sequences *i* and *j*.
- *L*: sequence length.
- *S*_*i*_[*k*], *S*_*j*_[*k*]: amino acids at position *k*.
- BLOSUM(*S*_*i*_[*k*], *S*_*j*_[*k*]): Raw BLOSUM62 log-odds score.
- *D*_*seq*_[*k*]: per-position distance (inverse likelihood).

#### 2.3.2 TCR V*β* Preference Distance

For each HLA, we calculate the V*β* gene preference *F* (Section 2.2). Difference between two vectors *F*_*i*_ and *F*_*j*_ are measured with cosine distance:

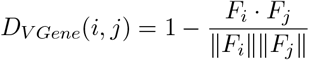

where:

- *F*_*i*_ and *F*_*j*_: V*β* preference vectors for HLAs *i* and *j*.
- *F*_*i*_ · *F*_*j*_: dot product.
- ∥*F*_*i*_∥ and ∥*F*_*j*_∥: vector magnitudes.

### 2.3.3 Peptide Motif Distance

For each HLA, aligned peptide sequences from the MHC Motif Atlas dataset (Tadros et al., 2023) are used to build a peptide motif matrix as a PWM. Distances between peptide motifs are measured with JS divergence, yielding a symmetric distance matrix.

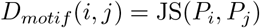

where:

- *P*_*i*_ and *P*_*j*_: PWMs for HLAs *i* and *j*.

The following HLAs are absent from the Atlas and excluded from peptide analysis: DPA1 *×* 01:03+DPB113:01, DQA101:02+DQB106:09, DQA101:03+DQB106:01, DQA101:04+DQB105:03, DQA101:05+DQB105:01, DQA103:03+DQB102:02, DQA103:03+DQB103:01, DQA104:01+DQB1 *×* 04:02, DQA105:05+DQB103:01.

### 2.3.4 Distance Correlations

To capture non-linear relationships among *D*_*seq*_, *D*_*V Gene*_, *D*_*motif*_, we use Spearman correlation with a stringent 1% False Discovery Rate (FDR) *N* = 1000 bootstrapping replicates to ensure robust correlation estimates.

Each HLA position is assigned a correlation coefficient and standard deviation for TCR*β* usage and peptide motifs, enabling identification of positions more strongly correlated with one or the other.

To avoid redundancy, correlations are computed only on the lower triangle of the distance matrices. For any pairs considered (*i, j*) where *i* < *j*:

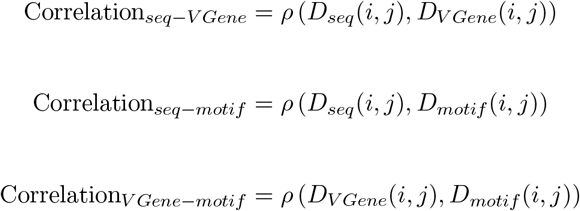

We also report the overall correlations from the full distance matrices.

### 2.4 HLA Position Importance Disentanglement

For each HLA position, Correlation_*seq-V Gene*_ and Correlation_*seq-motif*_ provide coefficients for TCR*β* usage and peptide motifs. Significant position are plottedin Figure 4C (per-locus plots and perposition values in Appendix E).

These positions are highlighted in their respective 3D structure for each locus (see Appendix E.3) and overall (see Figure 4D).

### 2.5 V*β* Promiscuity and HLA Restriction

To understand the breadth of HLA restriction for individual TCR V*β*s, we calculate the conditional probability *P* (HLA_*i*_ | V*β*_*j*_). While our previous analysis focused on HLA preferences (*P* (V*β* | HLA)), this “V-gene centric” view quantifies how specific or promiscuous a V*β* is across the HLA landscape.

We derive this by column-normalizing the co-occurrence matrix *C*:

- *C*_*ij*_: count of TCR V*β j* for HLA *i*.
- 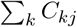 total count of V*β j* across all HLAs.
- Restriction probability:

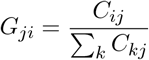

Each row *j* of matrix *G* represents the probability distribution of HLA alleles associated with a given V*β*. To identify V*β*s with shared HLA binding preferences, we perform hierarchical clustering on the columns using cosine distance.

Finally, we quantify the “promiscuity” of each V*β* using the Shannon entropy of its HLA distribution:

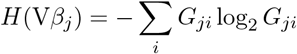

To compare restriction breadth across HLA loci with varying numbers of alleles (*N*), we calculated the *normalized entropy* (*H*_norm_ = *H/* ln *N*), which scales the measure between 0 and 1. We interpret V*β*s with low normalized entropy as “specialists” and those with high normalized entropy as “generalists.”

Differences in normalized entropy distributions between loci were assessed using pairwise Mann-Whitney U tests. To account for multiple comparisons across all locus pairs, p-values were adjusted using the Bonferroni correction.

## 3 Results

TCR recognition of pHLA complexes is primarily mediated by three complementarity-determining regions (CDRs). CDR1 and CDR2 are encoded by the V*β* gene, while the hypervariable CDR3 region is generated by V(D)J recombination (Davis and Bjorkman, 1988; Murphy and Weaver, 2017b). To better understand how germline-encoded TCR features and HLA polymorphisms shape pHLA recognition, we examine how HLA polymorphism correlates with TCR V*β* gene usage, focusing on CDR1 and CDR2 regions and distinguishing direct HLA effects from peptide-mediated influences. Our goal is to disentangle the contributions of direct TCR–HLA interactions from those arising indirectly through variation in peptide presentation.

Our analysis draws on large-scale repertoire sequencing data linking TCR*β*s with HLAs across a cohort of approximately 30,000 individuals (see Methods Section 2.1). V*β* gene usage patterns for common HLAs are derived from TCR*β*s statistically associated with each HLA and aggregated into vectors of normalized frequencies (Supplementary Figure S2), which we interpret as V*β* gene preferences of each HLA. To validate that these patterns are stable and not confounded by individual immune exposure, we compare repertoires from Cytomegalovirus (CMV) -positive and CMV-negative subjects, restricting analysis to HLA-associated TCRs observed in individuals carrying the relevant HLA allele. We use CMV exposure as a test case as it typically has a strong impact on the T cell repertoire (Klenerman and Oxenius, 2016) and can be inferred directly from the TCR*β* repertoire (Emerson et al., 2017; May et al., 2024). This makes CMV a strong test case: if a single dominant exposure perturbs V*β* usage patterns, we should detect it in CMV-positive subjects. As shown in Figure 2, V*β* usage is consistent between CMV-positive and CMV-negative subjects, supporting the conclusion that V*β* gene preferences reflect HLA-driven selection rather than exposure-dependent biases. Using the T-Detect COVID cohort (Zahid et al., 2025b) of approximately 30,000 individuals, we infer both CMV exposure status (Emerson et al., 2017) and HLA genotype (Zahid et al., 2025a). We focus on individuals carrying HLA-A02:01, comparing V*β* usage among CMV-positive (N = 5,449) and CMV-negative (N = 8,136) subjects. For each person, we quantify V*β* usage across all HLA-A02:01–associated TCR*β* sequences in their repertoire (45,737 unique sequences). We observe no significant difference in V*β* usage patterns between the CMV-positive and CMV-negative groups (see Figure 2). Restricting the analysis to other HLAs yields consistent results.

Although this analysis does not rule out contributions from other antigen exposures, it shows that even a strong immune stimulus like CMV does not measurably influence HLA-associated V*β* usage patterns. This supports the interpretation that the observed associations reflect generalizable, population-level trends.

Next, we analyzed how HLA polymorphism influences TCR*β* recognition and peptide binding. We represent HLA polymorphism using aligned sequences trimmed to the TCR-facing surface, including peptide-binding domains (Bradley, 2023). Peptide repertoires for each HLA are obtained from the MHC Motif Atlas (Tadros et al., 2023) and represented as Position Weight Matrices (PWMs), which capture amino acid frequencies across positions (Parker et al., 1994) (Supplementary Table S1). Then, we define three pairwise distance metrics between HLA alleles (e.g., *A02:01* vs. *A02:02*): (i) HLA sequence divergence, calculated from BLOSUM62-derived substitution scores (Henikoff and Henikoff, 1992), either summed across aligned residues to yield a global similarity score or evaluated at individual positions for residue-level analysis; (ii) TCR V*β* usage similarity, computed as the cosine distance between normalized V*β* gene frequency vectors; and (iii) peptide motif divergence, measured as Jensen-Shannon (JS) divergence between PWMs. This framework enables us to assess whether specific HLA polymorphisms are more strongly associated with the peptide repertoire or with the set of TCR*β*s likely to recognize a given HLA. A summary of the analysis workflow is shown in Figure 1.

**Figure 1.**
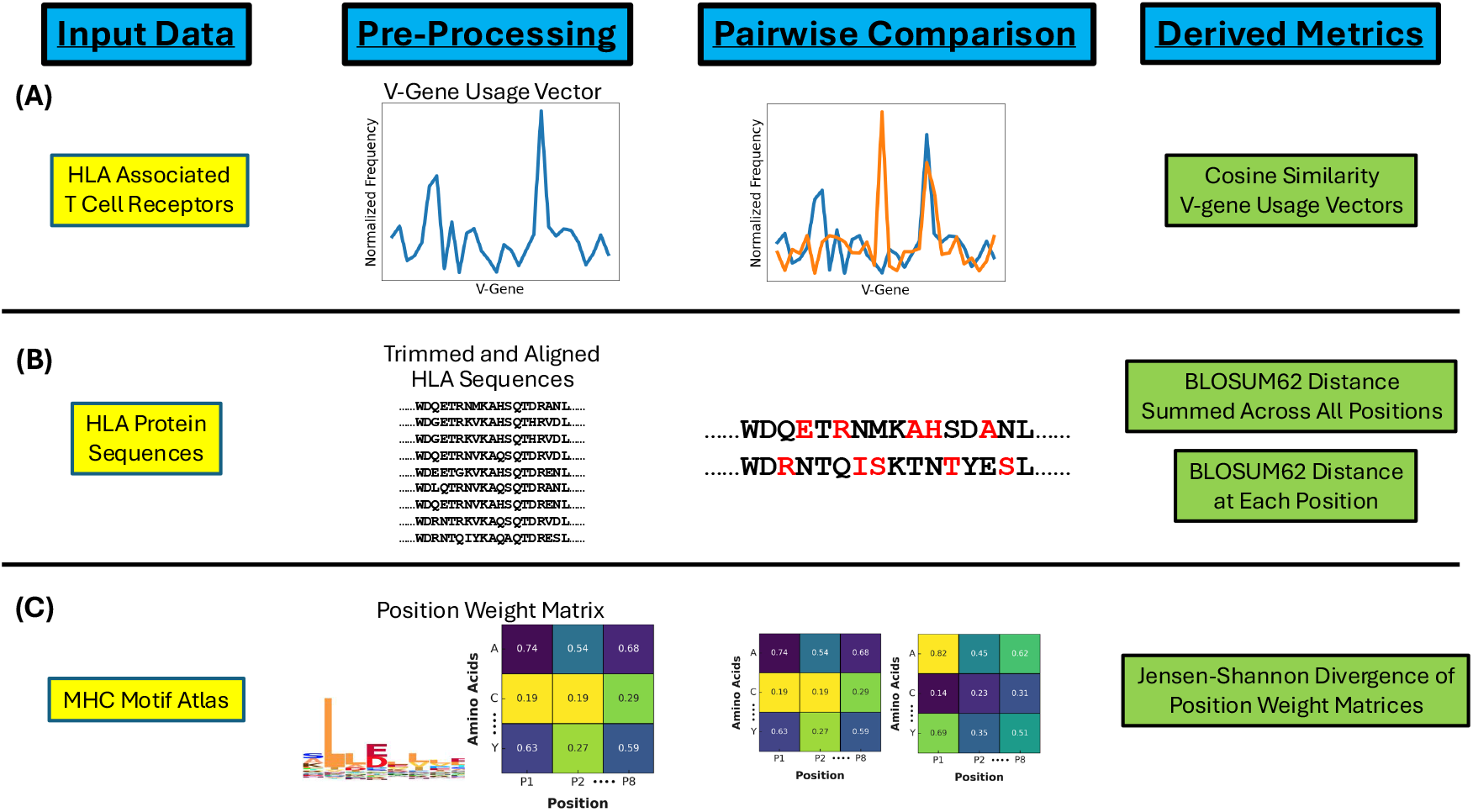
Summary of our analysis workflow. **(A)** Starting from sets of TCR*β*s associated with specific HLAs, we generate normalized V*β* gene usage vectors for each HLA. We quantify TCR V*β* usage similarity in a pairwise manner, calculating the cosine distance between the V*β* gene usage vectors of two HLAs. **(B)** Starting from HLA protein sequences, HLA sequences are trimmed to the TCR-binding surface and aligned via Multiple Sequence Alignment (MSA) (Bradley, 2023). We quantify both overall and position-dependent HLA polymorphism by comparing HLA sequences using the BLOSUM62 substitution matrix. **(C)** We use MHC Motif Atlas (Tadros et al., 2023) to generate Position Weight Matrices (PWMs) of the peptide repertoire for a given HLA. PWMs encode the same information as logo plots, indicating the frequency of amino acid use at any given position. Using PWMs, we calculate the Jensen-Shannon Divergence to quantify similarity of the peptide repertoire of any two HLAs.

**Figure 2.**
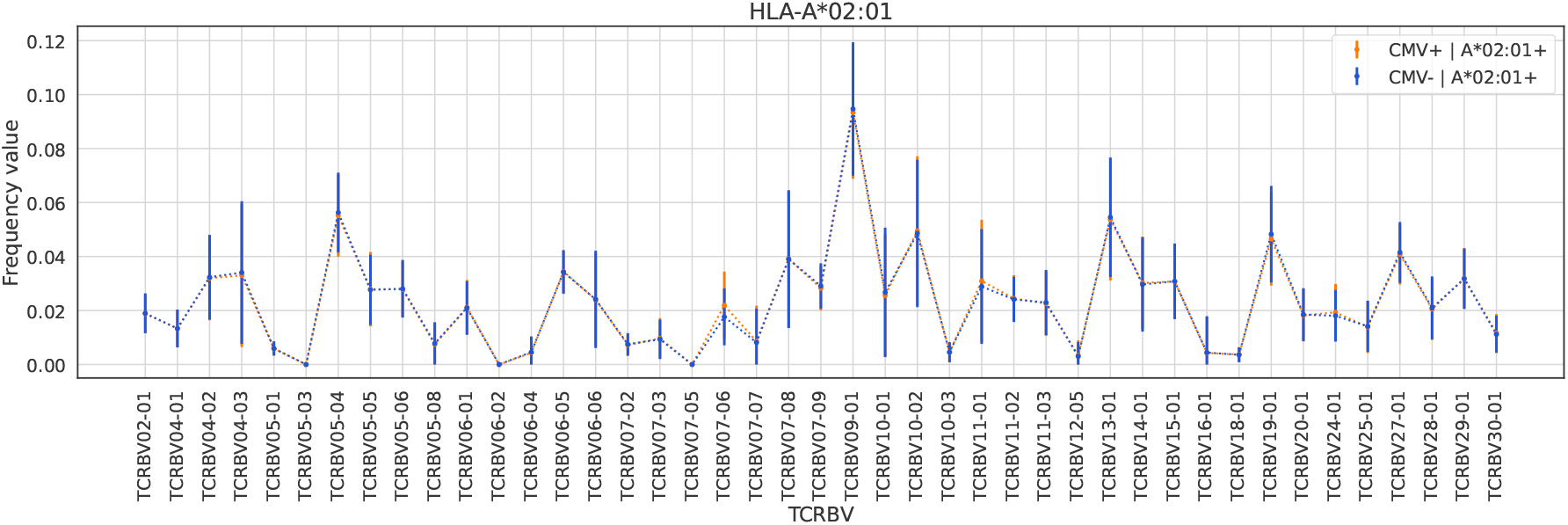
TCR V*β* usage patterns among HLA-A*02:01-associated sequences are unaffected by CMV exposure. Background-normalized V*β* gene usage across HLA-A*02:01-positive donors is shown for CMV-positive (orange) and CMV-negative (blue) cohorts. Error bars indicate the standard deviation across individuals within each group. Usage patterns are consistent between CMV-positive and CMV-negative subjects, suggesting that CMV exposure does not influence HLA-associated V*β* preferences.

Before examining how HLA sequence variation relates to similarity in V*β* gene usage across alleles, we first characterize the overall breadth of V*β* restriction exhibited by different HLA loci.

### 3.1 Variation in V*β* Preference Breadth Across HLA Loci

To assess how HLA polymorphism affects the breadth of the T cell repertoire, we quantified how “exclusive” the V*β* gene usage was for each HLA gene using Shannon entropy (*H*_norm_) on the conditional probability distribution *P* (HLA | V*β*). In the case of HLA-associated TCRs restricted to single V*β* gene, the entropy score would be close to 0. Conversely, uniform usage of all V*β* genes would yield a score closer to 1.

As shown in Supplementary Figure S18, there is a significant difference in restriction breadth across loci. Class II loci HLA-DR and HLA-DP displayed normalized entropy values approaching the theoretical maximum (*H*_norm_ ≈ 0.99), corresponding to a nearly uniform distribution of V*β* usage across alleles. In contrast, HLA-C exhibited a significantly lower mean entropy and higher variance compared to HLA-DR (see Supplementary Table S10). HLA-C shows the lowest entropy and highest variance across loci, indicating a high degree of restriction due to specific V*β* genes being associated with a limited subset of alleles. HLA-A, -B and -DQ display intermediate entropy profiles significantly distinct from both the HLA-C and HLA-DR/DP extremes (Supplementary Table S10)

These differences highlight substantial locus specific variation in the overall permissiveness of V*β* gene usage. With this baseline heterogeneity in mind, we next examine how differences in HLA sequence relate to similarity in V*β* gene preferences and peptide binding across alleles.

### 3.2 Global Correlations Across HLAs

To test whether HLA polymorphism systematically shapes TCR and/or peptide features, we analyze correlations between global HLA sequence similarity (summed across all aligned positions), V*β* gene usage and peptide motif similarity within each HLA class. We find that overall HLA sequence similarity correlates with both TCR V*β* gene preference (Figure 3A) and peptide motif similarity (Figure 3B; Table 1), even when using partial Spearman correlation to control for HLA locus structures. These results suggest that variation in HLA sequence alone is sufficient to predict meaningful trends in V*β* gene usage, highlighting the potential for sequence-based features to inform models of TCR–HLA specificity.

**Table 1:** Spearman *ρ* coefficients and bootstrap standard deviation (SD) of HLA sequence polymorphism measured with BLOSUM Distance, comparing TCR V*β* gene Preference with cosine distance and Peptide Motifs with JS Divergence for all loci and classes. A significance threshold of 0.01 was used and p-Values below this threshold are in bold. See details about TCR-HLA in Appendix Section C and HLA-Peptide in Appendix Section D.

| Class | Locus | # Alleles | HLA-TCR |  | HLA-Peptide |  |
| --- | --- | --- | --- | --- | --- | --- |
| | | | Spearman $\rho \pm \text{SD}$ | p-Value | Spearman $\rho \pm \text{SD}$ | p-Value |
| 1 | all | 34 | **0.30 ± 0.04** | **4.43E-13** | **0.25 ± 0.04** | **1.08E-09** |
| 2 | all | 57 | **0.60 ± 0.02** | **1.07E-155** | **0.52 ± 0.02** | **1.04E-74** |
| 1 | A | 12 | **0.35 ± 0.11** | **4.17E-03** | **0.57 ± 0.09** | **4.31E-07** |
| 1 | B | 17 | **0.29 ± 0.08** | **7.78E-04** | **0.33 ± 0.08** | **9.76E-05** |
| 1 | C | 5 | -0.18 ± 0.38 | 6.51E-01 | 0.57 ± 0.28 | 8.97E-02 |
| 2 | DP | 12 | 0.26 ± 0.12 | 3.22E-02 | **0.80 ± 0.06** | **2.22E-13** |
| 2 | DQ | 16 | **0.27 ± 0.09** | **3.19E-03** | **0.48 ± 0.15** | **8.31E-03** |
| 2 | DR | 29 | **0.48 ± 0.04** | **1.00E-24** | **0.36 ± 0.05** | **5.25E-14** |

**Figure 3.**
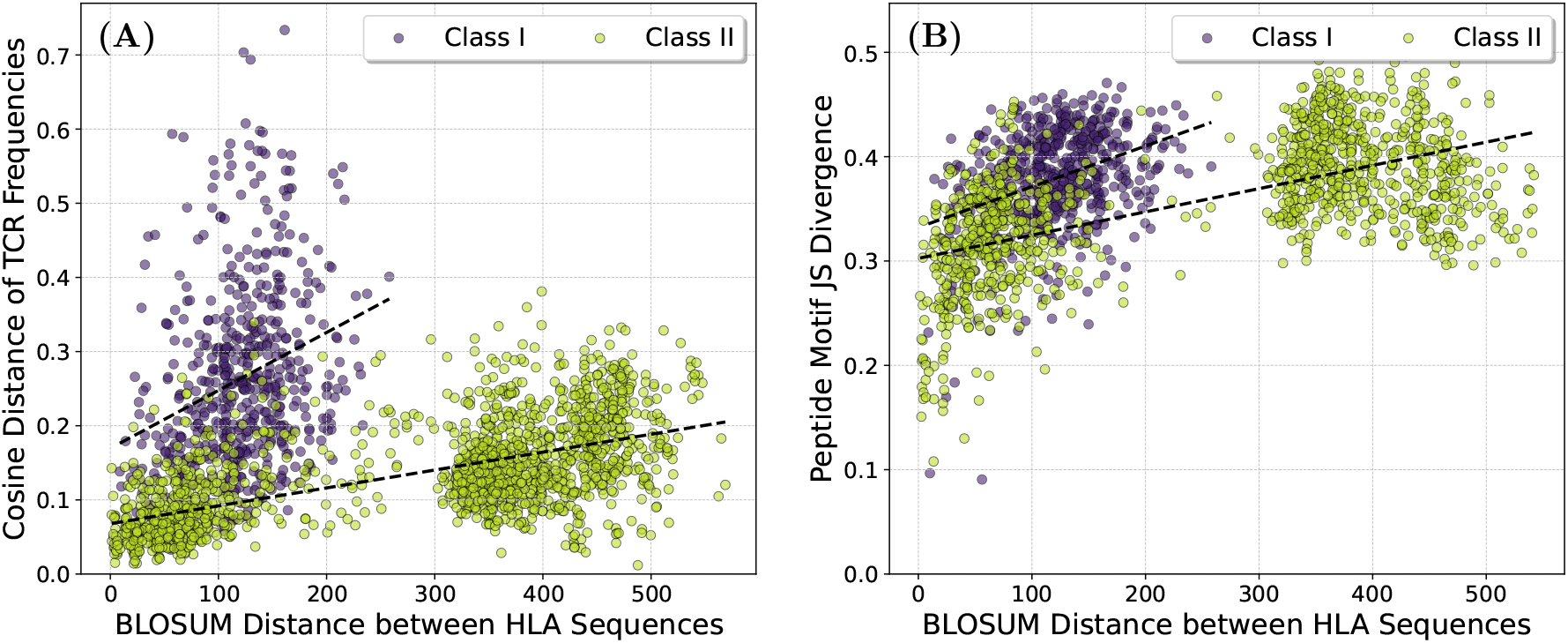
A pairwise comparison showing the relationship between HLA polymorphism, V*β* gene usage and the peptide repertoire. **(A)** TCR V*β* usage similarity plotted against HLA polymorphism. HLAs with similar sequences tend to have similar V*β* gene preferences. Dotted lines illustrate correlations which we quantify using partial Spearman’s *ρ* (class I: *ρ* 0.30 ± 0.04, class II: *ρ* 0.60 ± 0.02). **(B)** Peptide repertoire similarity plotted against HLA polymorphism. HLAs with similar sequences tend to have similar peptide repertoires. Dotted lines illustrate correlations which we quantify using partial Spearman’s *ρ* (class I: *ρ* 0.25 ± 0.04, class II: *ρ* 0.52 ± 0.02). See Table 1 for per locus Spearman’s *ρ* and p values and Appendix Section C for TCR-HLA and Appendix Section D.1 for HLA-peptide.

At the locus level, HLA sequence similarity correlates with both V*β* gene and peptide features for loci A, B, DQ and DR (see Table 1 and Figure 4). DR shows the strongest association with V*β* gene preference, while DP shows the strongest correlation with peptide motifs but not with V*β* gene features. Locus C does not show significant correlations in any analysis, likely due to its lower surface expression and resulting in limited statistical power (Neefjes and Ploegh, 1988). Notably, we observe that V*β* gene usage frequencies are more variable among Class I alleles compared to Class II, which exhibit more uniform usage patterns across alleles (Supplementary Figure S3).. These findings indicate that the relationship between HLA polymorphism, TCR*β* engagement and peptide binding is both locus- and class-dependent, with distinct correlation patterns emerging across class I and class II molecules.

**Figure 4.**
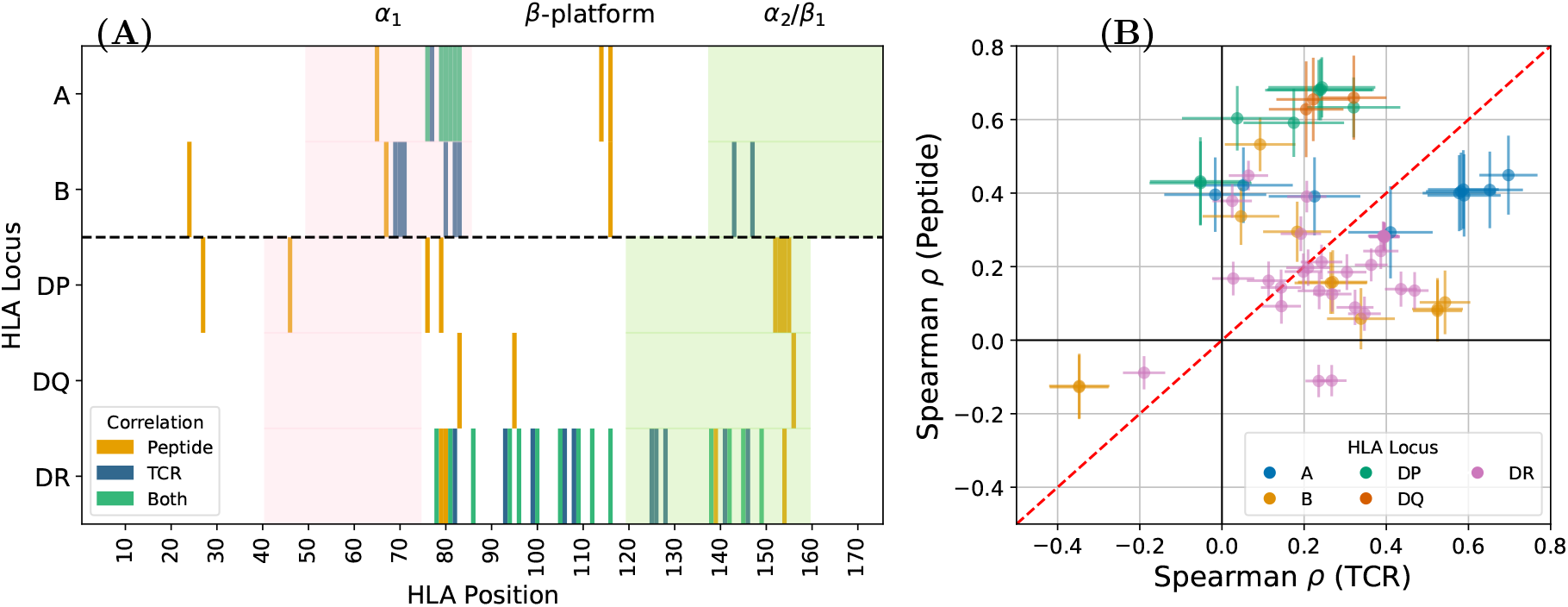
Disentangling HLA positions correlated with V*β* gene usage vectors, the peptide repertoire or both. **(A)** Across all loci, HLA polymorphic positions significantly correlated (p-value ≤ 0.01) with similarity in V*β* gene usage (blue), peptide repertoire similarity (yellow) and both (green). Shaded are the rough positions of the HLA binding groove helices. To allow positional comparison, for Class II we combined the *α*1 and *β*1 sequence. In light red is the *α*1 helix region (closest to TCR*β*). In light green is the *α*2 helix (or *β*1 for Class II) (closest to TCR*α*). See also per-locus plots in Appendix C (TCR*β*) and D (peptide). **(B)** Scatter plot of Spearman’s *ρ* values and bootstrapped standard deviation for the correlation of HLA polymorphism with peptide motif similarity versus its correlation with V*β* gene usage similarity. Only HLA positions where at least one significant correlation is derived are plotted. See per-locus plots and per-position values in Appendix E.

To further probe the structural basis of these patterns, we restrict our analysis to HLA residues known to contact either the peptide or the TCR, based on solved pHLA–TCR structures. This restriction strengthens the observed correlations (see Supplementary Figure S1), suggesting the correlation is driven by a subset of positions and thus motivating a focused, residue-level investigation of TCR–HLA specificity.

### 3.3 Residue-Level Correlation Analysis

We hypothesize that polymorphic HLA residues influence TCR recognition through distinct mechanisms, with some acting directly on TCR engagement while others act indirectly via influence on peptide binding. To test this, we analyze residue-specific HLA polymorphisms, which are largely concentrated in the peptide-binding groove, while more distal regions remain conserved (Supplementary Figure S5). This pattern of polymorphism further suggests that only a subset of residues drive correlations with V*β* gene usage and peptide motifs. We quantify each residue’s contribution by correlating per-residue polymorphism with both V*β* gene preference and peptide motif similarity (see Supplementary Figures S8 and S11). This analysis identifies HLA residues statistically associated with TCR recognition, peptide binding or both.

We visualize the spatial distribution of residues significantly associated with V*β* gene usage or peptide motifs across class I and class II loci (Figure 4; see Supplementary Sections B–D for details). Peptide-associated residues are broadly distributed across the binding groove, though in HLA-A they cluster near the *α*1, closest to the TCR*β*. Residues associated with V*β* gene usage appear primarily in loci A, B and DR, clustering near the TCR-facing helices in class I and spanning the polymorphic DRB chain in class II. As expected, significant positions in DR are restricted to the polymorphic DRB chain, as the invariant DRA chain does not contribute variation in our polymorphism-based analysis. These findings suggest that residues correlated with peptide and TCR features occupy overlapping but structurally distinct regions, with class-specific patterns across loci.

To compare the relative influence of each HLA position on TCR V*β* usage and peptide presentation, we compute residue-specific Spearman correlation coefficients for both signals. For each polymorphic residue, we calculate its correlation with V*β* gene preference and peptide motif similarity. We then plot all positions with at least one significant correlation in Figure 4B. Points above the identity line indicate stronger correlations with peptide motifs, while those below indicate stronger correlations with V*β* gene usage. Positions in loci A and B are fairly balanced around the line, suggesting a mix of peptide- and TCR-associated effects. In contrast, most significant positions in loci DP and DQ are peptide-associated, while positions in DR cluster near the identity line with a modest bias toward TCR association. These results highlight locus-specific differences in how HLA residues influence TCR and peptide features, with distinct correlation patterns across class I and class II loci.

Surprisingly, several positions exhibit negative correlations, indicating that sequence differences at these sites are associated with greater similarity in peptide motifs or V*β* gene usage (see Supplementary Table S4 to Table S8). Although the reasons underlying this effect remain unclear, we hypothesize that such positions may reflect compensatory or co-evolutionary structural effects, where variation at one site may alter the HLA conformation in a way that preserves peptide presentation and/or TCR binding despite sequence divergence. Further structural and mutational studies are needed to test this hypothesis and clarify how these residues contribute to pHLA recognition.

### 3.4 Structural Distribution of Significant Positions

We visualize the significant positions on the 3D structure of an HLA molecule (Figure 5). Most fall within the peptide-binding groove, the most polymorphic region (Supplementary Figure S5). Residues more strongly correlated with peptide motifs (yellow) are primarily located on the *β*-sheets beneath the peptide and on the peptide-facing *α*-helices. In contrast, residues associated with TCR*β* features or with both TCR*β* and peptide features are predominantly found on the *α*-helices. In class I HLAs, these TCR- and dual-associated positions cluster near the region of the *α*1 helix (closest to the TCR*β*). In class II HLAs, they are more broadly distributed across the binding groove, particularly along the polymorphic DRB chain, consistent with the invariant nature of the DRA chain.

**Figure 5.**
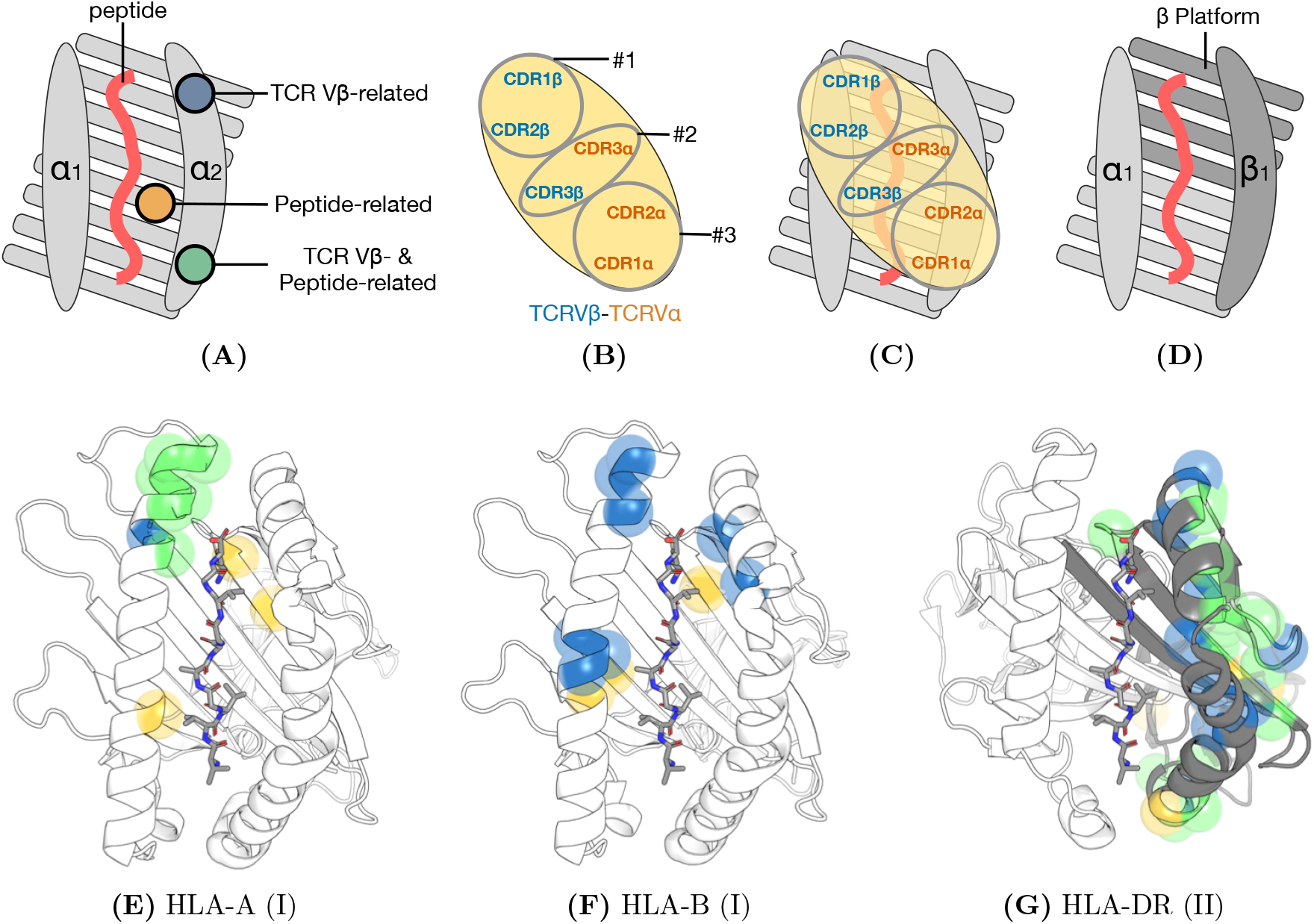
Structural logic of TCR–pHLA engagement and distribution of significant residues. **(A)** Schematic of the peptide-binding groove in Class I HLAs (*α*1 and *α*2 helices), illustrating three categories of polymorphic residues: those related to TCR V*β* (blue), peptide (yellow) or both (green). **(B)** Schematic of the TCR binding interface (footprint), showing the arrangement of CDR loops categorized into three functional ‘topes’ (Cohn et al., 2019): Tope #1 (TCR*β* CDR1/2) and Tope #3 (TCR*α* CDR1/2) engage the HLA helices, while Tope #2 (hypervariable CDR3s) primarily engages the peptide. The TCR V*β* domain (blue text, corresponding to Tope #1) typically contacts the *α*1 helix in the canonical diagonal orientation. **(C)** Overlay of the TCR footprint onto the Class I binding groove, illustrating the proximity of V*β* CDRs to the *α*1 helix. **(D)** Schematic of the Class II binding groove, formed by the *α*1 and *β*1 chains. Note that unlike Class I, for Class II HLAs, the *β* platform is encoded by the *α* chain (light grey) and *β* chain (dark grey). **(E–G)** Correlated positions highlighted on HLA-A (E), HLA-B (F) and HLA-DR (G). Residues are colored by their statistical association: TCR V*β* (blue), peptide (yellow) or both (green). These positions are identified in Fig. 4 (See details in Supplementary Section D).

Together, these analyses show that HLA polymorphisms shape TCR recognition and peptide presentation by partially overlapping but distinct sets of HLA polymorphic contacts. Some residues directly influence TCR engagement, others act indirectly via peptide binding and a subset contribute to both. The balance of these effects varies across loci, with clear differences between class I and class II HLAs. This residue-level map of functional variation offers a valuable foundation for refining models of TCR–pHLA specificity and guiding future structural and computational work in immunogenetics.

## 4 Discussion

Our analysis leverages a large-scale statistical framework, using 30,000 TCR*β* repertoires to associate specific TCR*β*s with individual HLAs. By averaging across thousands of individuals and diverse but commonly shared antigenic exposures, we derive normalized V*β* gene preferences for each HLA, enabling a robust, population-level assessment of TCR*β*-HLA preferences. This approach yields sets of TCR*β*s that are statistically enriched for HLA specificity, providing statistical power to link TCR*β*s to their restricting HLAs despite the polygenic nature of HLA phenotypes.

To understand how HLA polymorphism shapes peptide binding and TCR engagement, we analyze correlations between HLA sequence similarity, peptide motifs and V*β* gene usage across the cohort. Our goal is to distinguish direct TCR–HLA associations from peptide-mediated effects. We find that HLA sequence similarity correlates with both V*β* gene usage and peptide motifs. These associations map to specific polymorphic residues linked to either V*β* preference, peptide features or both. Structurally, key positions cluster around the *α*1 helix in class I and the *β*1 helix in class II, with particularly strong signals in the DR locus. These results suggest that TCR–HLA specificity is shaped by a combination of germline-encoded contacts and indirect peptide-driven effects, including potential allosteric modulation of pHLA conformation. Together, these patterns highlight distinct structural contributions to recognition across HLA classes. Notably, the clustering of TCR-associated residues on the Class I *α*1 helix structurally confirms the germline contact maps proposed by Marrack et al. (2008).

A longstanding question in immunology is whether TCR–pHLA specificity is governed more by inherited structural constraints or by developmental selection, reflecting broader questions about how evolutionary forces (Feng et al., 2007; Marrack et al., 2008; Christopher Garcia et al., 2009; Yin et al., 2012) and thymic processes (Van Laethem et al., 2007, 2012; Lu et al., 2019) shape the TCR repertoire. Evidence for intrinsic specificity includes recognition of pHLAs by randomly generated T cells (Krovi et al., 2019), QTL associations between HLAs and TCR V*β* usage (Sharon et al., 2016; Gao et al., 2019) and alloreactivity (Felix and Allen, 2007), where TCRs selected on self-HLAs cross-react with non-self HLAs. Our findings support this view. Namely, polymorphic residues, especially in the *α*1 helix of class I and the *β*1 helix of class II, correlate with V*β* gene preference but not peptide motifs, consistent with germline-encoded structural bias and allele-specific recognition (Cohn et al., 2019). We also identify residues associated only with peptide motifs and those associated with both peptide motifs and V*β* genes, indicating multiple mechanisms through which HLA polymorphism shapes TCR engagement. A parsimonious interpretation is that TCR–HLA interactions reflect both co-evolution and selection (Rangarajan and Mariuzza, 2014; La Gruta et al., 2018), with germline contacts **priming** initial binding and thymic selection, subject to these germline constraints, **sculpting** the final repertoire. This layered model helps reconcile competing hypotheses and offers a unified framework for understanding repertoire architecture.

Our analysis suggests that V*β* restriction follows a continuum rather than a simple Class I versus Class II split. HLA-DR and HLA-DP appear to be “generalists” with high entropy and broad compatibility. On the other hand, HLA-C seems to act as a “specialist” with lower entropy compared to other HLAs. This complements Gao et al. (2019)’s observation that HLA-C has fewer TCR associations than other Class I loci. While they attribute this to low surface expression, our data implies that these associations are more specific for HLA-C than for other loci. Therefore, the weak HLA-C signal could result from a combination of low abundance and structural selectivity. This divergence in part reflects the functional needs of CD8+ T cells to have precise recognition to avoid autoimmunity, whereas CD4+ T cells benefit from broader scanning capabilities.

Several findings raise compelling questions for future investigation. In class I HLAs, V*β* gene associations cluster near the *α*1 helix, which directly contacts the TCR*β* chain. In contrast, HLA-DR associations are exclusively driven by the polymorphic *β* chain (DRB), since the *α* chain (DRA) remains constant. This provides a residue-level mechanism for the *HLA-DRB1* expression QTLs reported by Sharon et al. (2016). Given that in Class II HLAs, the TCR*β* primarily contacts the *α*1 helix, it suggests that DRB polymorphisms may influence TCR binding indirectly, for example by changing pHLA conformation or stability, rather than through direct contact with the V*β* domain. The absence of similar correlations in HLA-DQ and -DP, where both chains are polymorphic, underscores the unique role of the invariant DRA chain in shaping recognition. Additionally, we identify HLA positions with negative correlations to TCR*β* or peptide similarity, potentially reflecting compensatory or co-evolutionary effects that preserve recognition despite sequence divergence. These patterns suggest novel structural mechanisms of immune recognition and point to the value of follow-up studies using mutagenesis, structural modeling or biophysical assays.

Despite the statistical power and scale of our framework, several limitations should guide interpretation. Our analysis focuses exclusively on the TCR*β* chain and does not account for potential contributions from the TCR*α* chain, which also engages the pHLA surface and may exhibit its own germline biases. Likewise, our emphasis on HLA polymorphism may overlook conserved germline-encoded residues that play critical roles in TCR recognition. While we focus on V*β* gene–encoded CDR1 and CDR2 regions, we do not account for the hypervariable CDR3 loop, which is central to peptide recognition and also influences HLA binding. In addition, peptide repertoires used to define motif similarity are drawn from public datasets often biased toward disease-associated antigens, potentially limiting their generalizability. Furthermore, some observed correlations may arise from linkage disequilibrium between HLA alleles, rather than from direct structural contacts. Conversely, the absence of correlation does not imply a lack of interaction but may reflect limited power, resolution or data sparsity. Structural modeling and molecular dynamics simulations could help resolve these ambiguities by assessing the functional impact of specific residues. Finally, as high-throughput technologies improve, future studies incorporating paired TCR*αβ* repertoires and more representative peptide libraries will be essential to validate and expand these insights.

A key assumption of our analysis is that the structural associations identified in the public repertoire reflect fundamental biophysical constraints applicable to all TCRs. Under this assumption, the V*β*–HLA associations we identify provide important constraints for modeling TCR-pHLA interactions. The *P* (*V β* | HLA) and *P* (HLA | *V β*) matrices provide simple priors that can bias a model toward germline-consistent TCRs when assigning HLAs or ranking candidate receptors. This is particularly useful in low-data settings (e.g. rare HLAs or small clinical cohorts), where these priors could help prevent models from overfitting to noise. These priors can also be incorporated into benchmarking via a simple prior-weighted accuracy metric.

Our findings support a model of TCR–HLA specificity shaped by both inherited constraints and adaptive processes. Germline-encoded biases appear to define the baseline potential for V*β* engagement with HLA, while thymic selection and immune experience further sculpt the repertoire. By leveraging tens of thousands of repertoires to derive V*β* usage patterns and mapping these onto HLA sequence and structure, we perform a residue-level analysis that provides insights into the molecular basis of TCR–HLA interactions. Our approach demonstrates the power of population-scale, structure-aware statistical analysis to uncover fundamental principles of immune recognition. These insights offer not only a clearer understanding of repertoire formation but also a scalable and interpretable framework to guide future efforts in TCR modeling, repertoire engineering and immunotherapy design.

## Supporting information

V gene data (with background)

## Author Contributions

LVC, MTN, HCH, and HJZ were involved in the writing of the paper. LVC worked on the analyses and code. LVC, MTN, HJZ created the visualizations. LP adviced on code structure. JG, HJZ, MTN, and HCH mentored LVC throughout the stages of the work. All authors provided feedback on the drafts of the paper.

## Acknowledgments

We thank Haiyin Chen for her persistent encouragement to pursue this problem, which ultimately motivated the work presented here.

## Supplemental Data

Supplementary Material should be uploaded separately on submission, if there are Supplementary Figures, please include the caption in the same file as the figure. LaTeX Supplementary Material templates can be found in the Frontiers LaTeX folder.

## Data Availability Statement

All data and code are available at https://github.com/microsoft/tcr-mhc.

