## Supplementary Materials for "HLA Alleles Imprint Distinct Biases in the Usage Preferences of TCR Vβ segments"

#### S1 Supplementary information

#### A Pseudo-sequence correlation

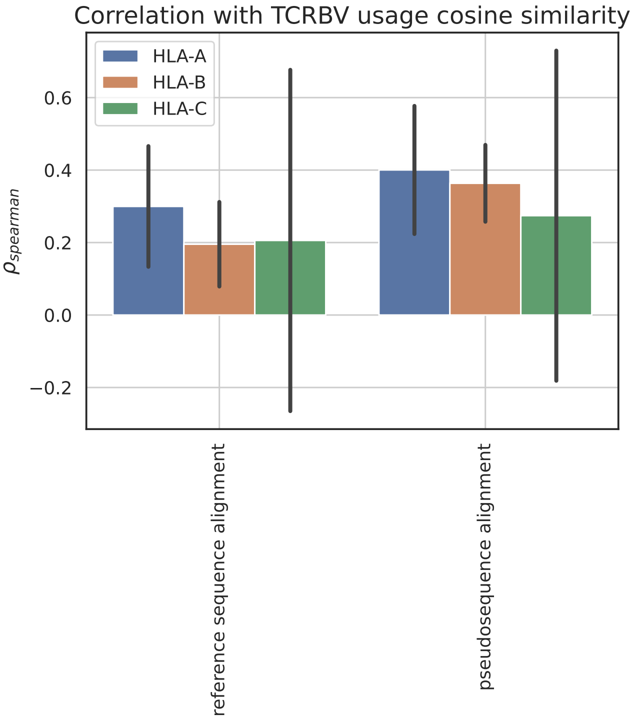


Figure S1: Spearman correlation coefficients between V$\beta$ usage similarity and HLA sequence similarity, shown separately for each class I locus (HLA-A, -B, -C). For each pair of HLAs, sequence similarity is computed in two ways: (1) by summing BLOSUM62 substitution scores across the full aligned HLA sequence and (2) by summing over a pseudosequence comprising only residues within 4 Å of the TCR in solved pHLA–TCR structures. Error bars represent the standard deviation across 1000 bootstraps, each selecting 1/2 of the alleles in each locus. Correlations based on the pseudosequence are consistently stronger across all three loci, suggesting that residues in direct contact with the TCR better explain V$\beta$ usage similarity.

We examine solved crystal structures to identify specific HLA residues that lie within 4 Å of the TCR, hypothesizing that these contact residues should exhibit stronger correlations with V$\beta$ gene usage. Due to limited structural data for class II HLAs, we restrict our analysis to class I loci. As shown in Figure S1, across all three class I loci, the correlation between V$\beta$ gene usage and the BLOSUM distance summed over the contact pseudosequence is stronger than when the score is summed over the full HLA sequence.

### B Exploratory Data Analysis

#### B.1 Data Overview

In addition to our internal dataset, we collated HLA sequence data from TCRDock (Bradley, 2023) and peptide data from the MHC Motif Atlas (Tadros et al., 2023). In Table S1, we summarize the number of data points for each HLA loci.

| Locus | Class | Number of HLAs | Number of TCRs | Number of Peptides |
|---|---|---|---|---|
| A | I | 12 | 256579 | 42374 |
| B | I | 17 | 416630 | 60843 |
| C | I | 5 | 88843 | 11690 |
| DP | II | 12 | 453742 | 18650 |
| DQ | II | 16 | 407645 | 8251 |
| DR | II | 29 | 1204418 | 66270 |

Table S1: Number of HLAs, TCRs and unique peptides per locus and class from TCRDock (Bradley, 2023) and MHC Motif Atlas (Tadros et al., 2023).

#### B.2 TCR$\beta$ preferences by HLA Locus

In Figure S2, we plot the TCR V$\beta$ preferences of each TCR for each HLA locus to examine intra-locus variability.

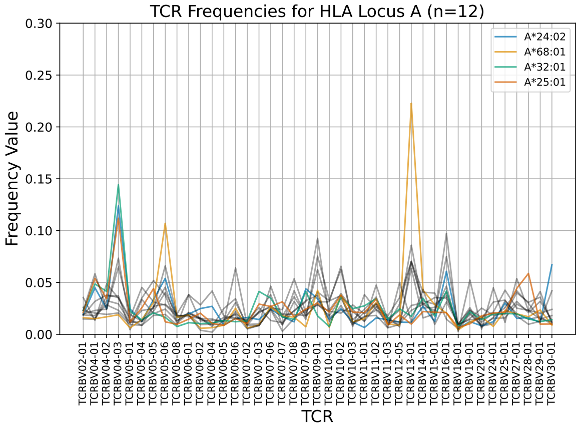


(a) Locus A

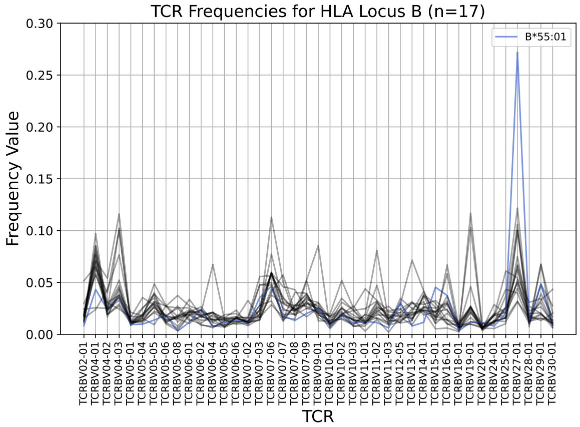


(b) Locus B

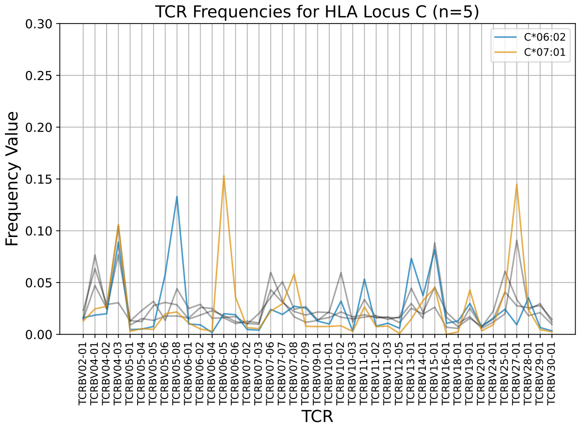


(c) Locus C

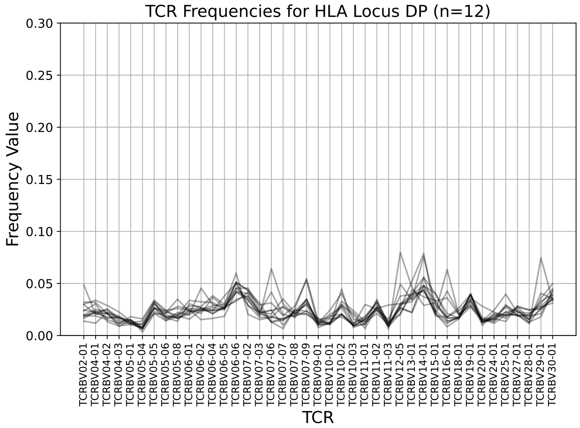


(d) Locus DP

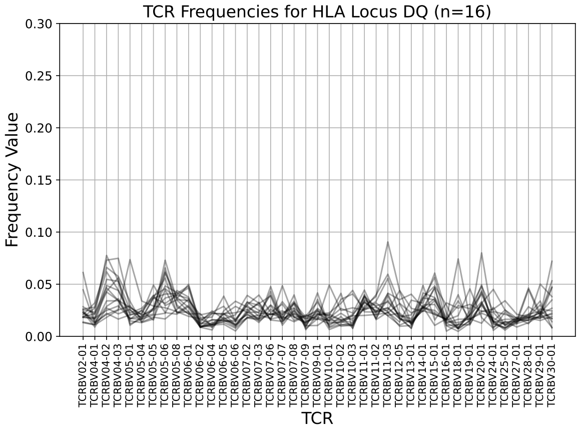


(e) Locus DQ

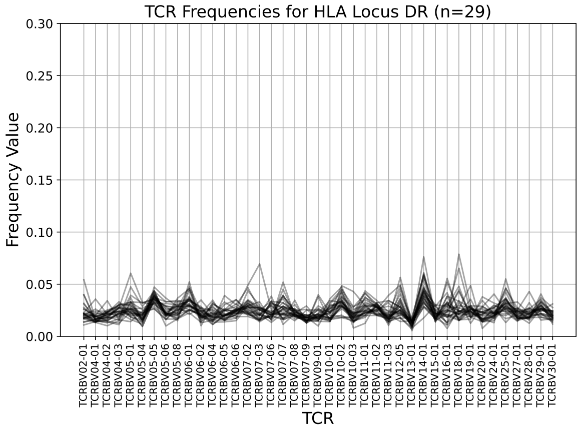


(f) Locus DR

Figure S2: Background-normalized TCR$\beta$ frequency values for each locus. Each line represents the values for one HLA.

#### B.3 Variance of V$\beta$ Usage Across HLA Classes

To examine the variability of V$\beta$ gene usage within each HLA class, we computed the variance of normalized TCR frequencies for each V$\beta$ gene across alleles of the same class. As shown in Figure S3, Class I HLAs (blue) exhibit a significantly broader distribution of variances compared to Class II (orange), which are tightly clustered near zero.

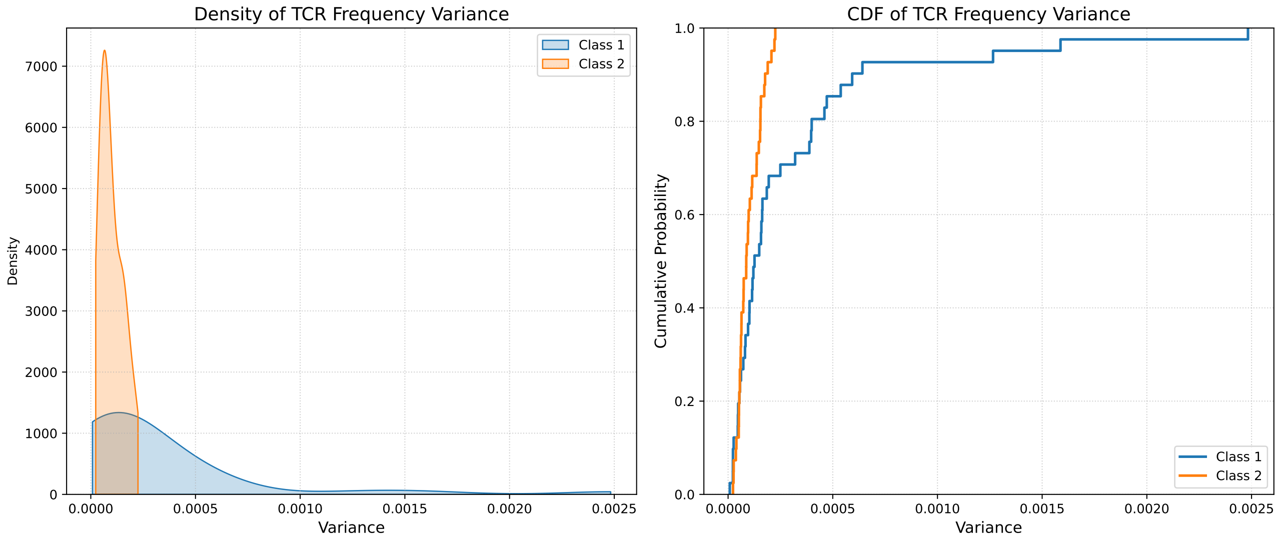


Figure S3: Distribution of TCR Frequency Variance by HLA Class. Left: Density plot (KDE) showing the distribution of variances for Class I (blue) and Class II (orange). Right: Cumulative Distribution Function (CDF) of the same data.

#### B.4 Overview of Distance Values

In Figure S4 we visualize the pairwise distance values of HLA sequence (BLOSUM Distance), TCR V$\beta$ preferences (Cosine Distance) and peptide motif Position Weight Matrix (JS Divergence). On the left we plot the histogram, while on the right we plot the cumulative distribution.

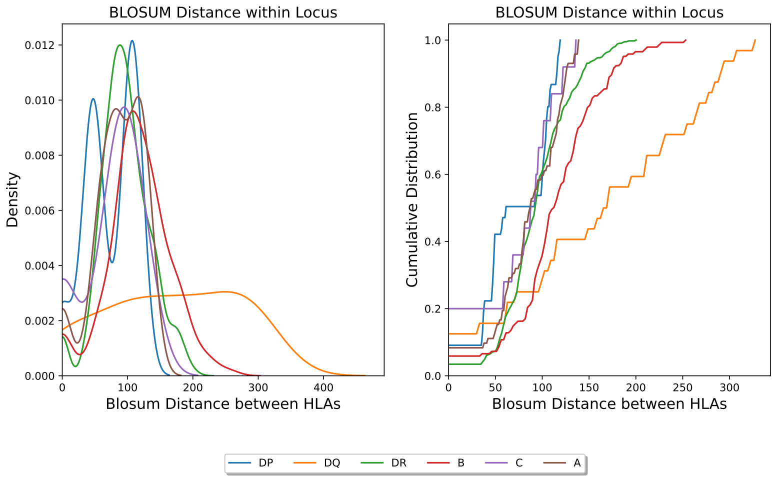


(a) $D_{seq}$: BLOSUM Distance distributions

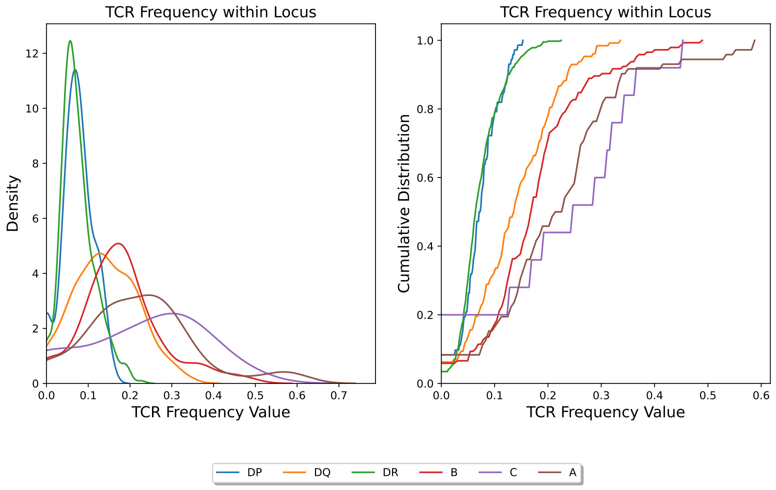


(b) $D_{VGene}$: TCR$\beta$ preferences distributions

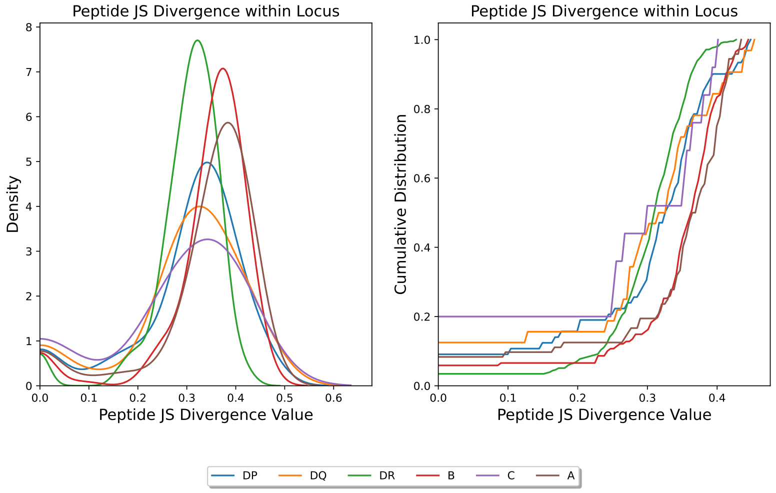


(c) $D_{motif}$: JS Divergence between Peptide Motifs

Figure S4: Histograms (left) and Cumulative Plot (right) of the distance matrices of HLA sequences (a), TCR$\beta$ preferences (b) and Peptide Motifs (c)

#### B.5 Overview of HLA Sequence Polymorphism on Structure

In Figure S5, we plot polymorphic positions for each HLA locus. After a sequence alignment using Clustal $\omega$(Sievers et al., 2011), we quantify polymorphism as the number of times a position has a different residue divided by the total number of sequences for that locus. The intensity of the red color corresponds to increased diversity for that position.

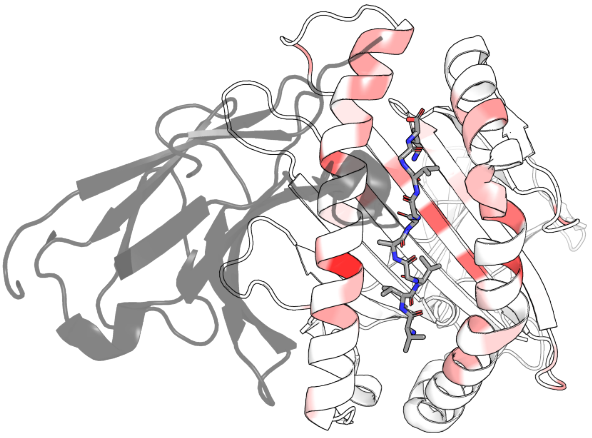

(a) Locus A

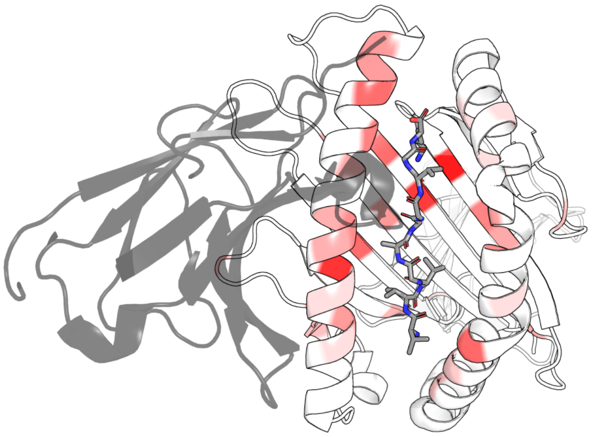

(b) Locus B

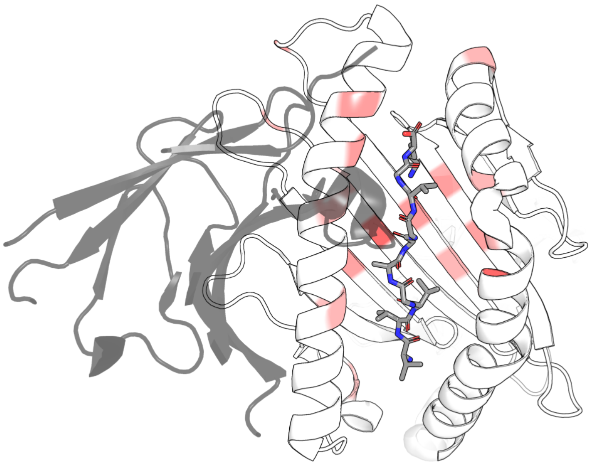

(c) Locus C

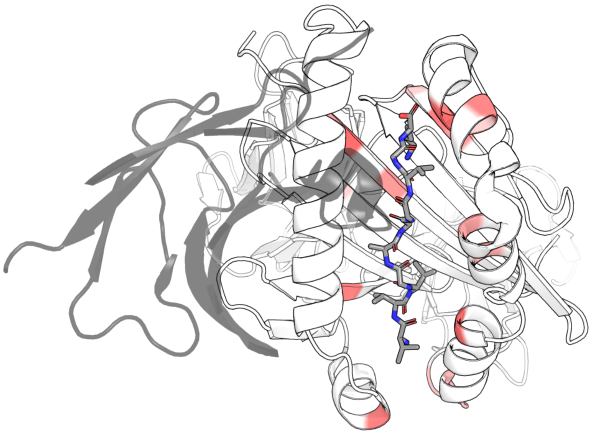

(d) Locus DP

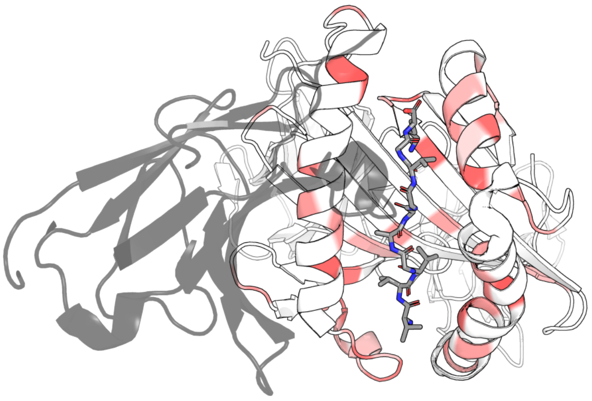

(e) Locus DQ

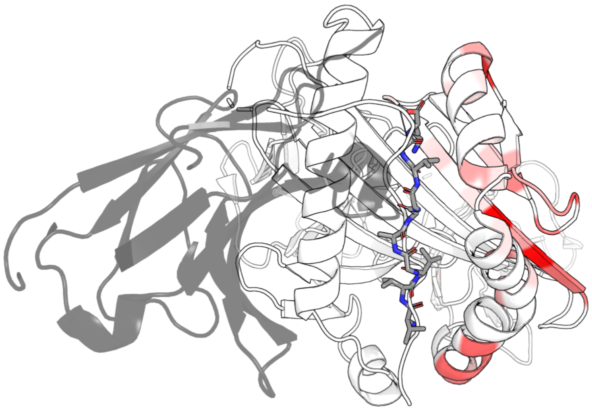

(f) Locus DR

Figure S5: Sequence polymorphism for each locus as measured by entropy. In grey the TCR$\beta$ contacting the $\alpha_1$ helix.

### C HLA Sequence Polymorphism and TCR$\beta$ Distance correlation

In this section we quantify the relationship between HLA polymorphism and changes in TCR V$\beta$ preferences using Spearman correlations. We break this down into an overall correlation (Section C.1), per-locus correlation (Section C.2) and per-locus and position in the HLA sequence (Section C.3) to identify the main drivers of the patterns.

#### C.1 Overall

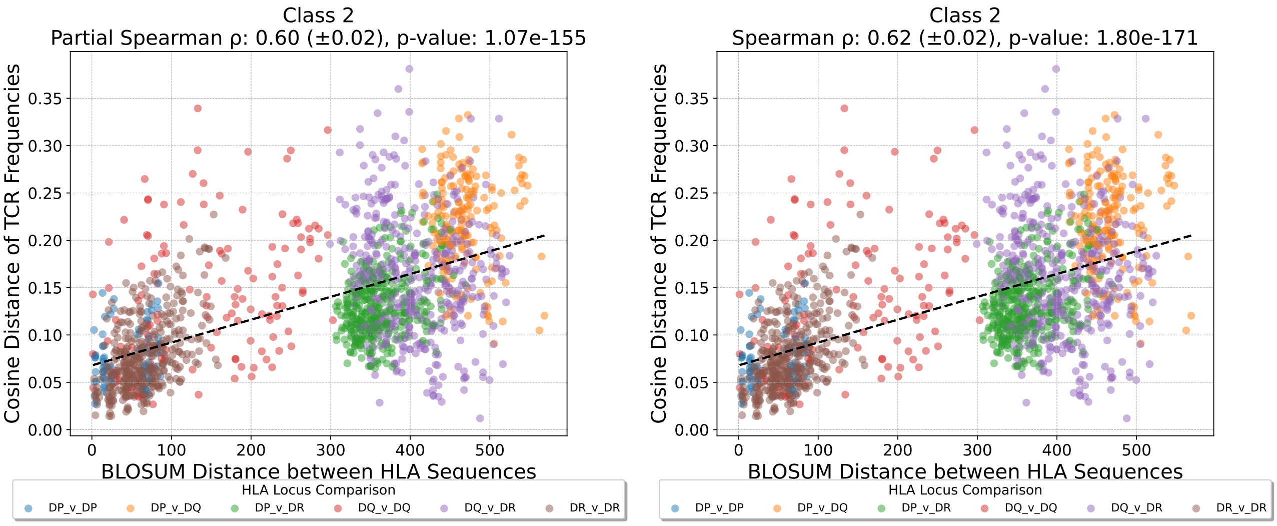


(a) HLA class I

(b) HLA class II

Figure S6: Relationship between HLA sequence polymorphism measured with BLOSUM Distance and TCR V$\beta$ Preference measured with cosine distance. The analysis is performed per HLA class and locus. See Table S2 for individual Spearman's $\rho$ values and Figure S7 for plots per locus.

| **Class** | **Locus** | **Spearman $\rho$ (Bootstrap SD)** | **p-Value** |
|---|---|---|---|
| 1 | all | 0.30 (0.04) | **4.43E-13** |
| 2 | all | 0.60 (0.02) | **1.07E-155** |
| 1 | A | 0.35 (0.11) | **4.17E-03** |
| 1 | B | 0.29 (0.08) | **7.78E-04** |
| 1 | C | -0.18 (0.38) | 6.51E-01 |
| 2 | DP | 0.26 (0.12) | 3.22E-02 |
| 2 | DQ | 0.27 (0.09) | **3.19E-03** |
| 2 | DR | 0.48 (0.04) | **1.00E-24** |

Table S2: Spearman $\rho$ coefficients and bootstrap standard deviation (SD) of HLA sequence polymorphism measured with BLOSUM Distance and TCR V$\beta$ Preference measured with cosine distance for all loci and classes, using a significance threshold of 0.01. The p-Values of the tests that pass the threshold are in bold. See Figure S7 for a per locus plot.

#### C.2 Per Locus

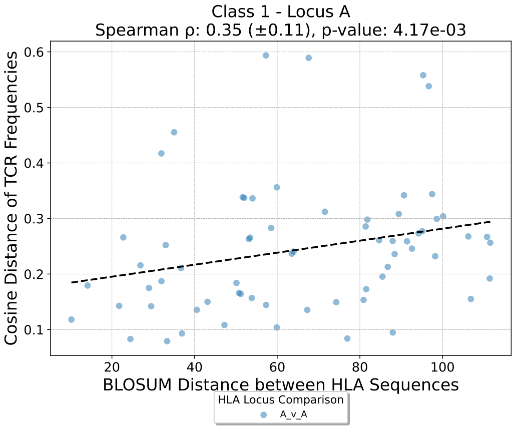


(a) Locus A

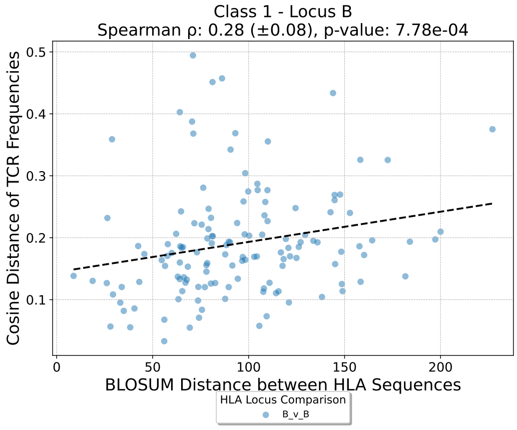


(b) Locus B

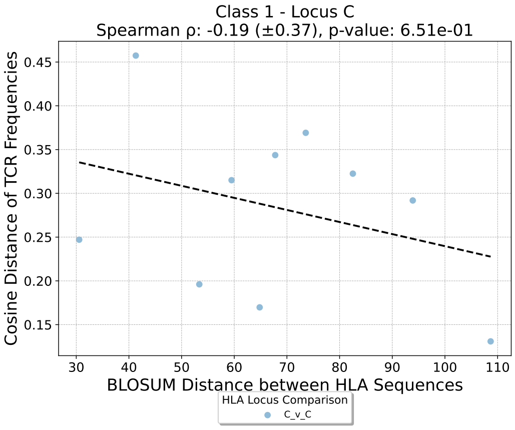


(c) Locus C

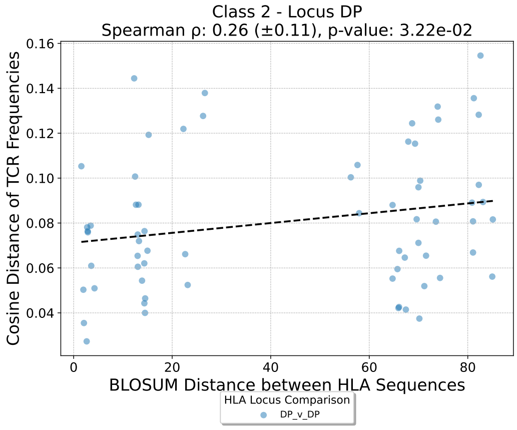


(d) Locus DP

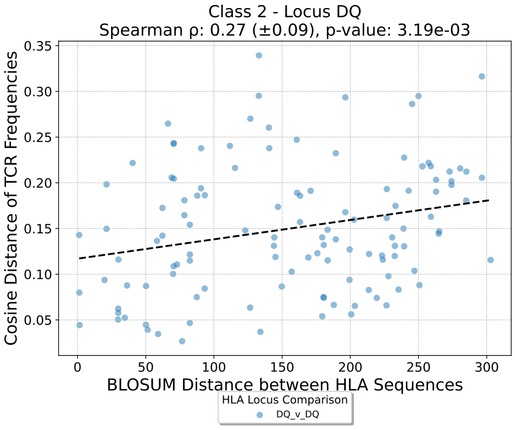


(e) Locus DQ

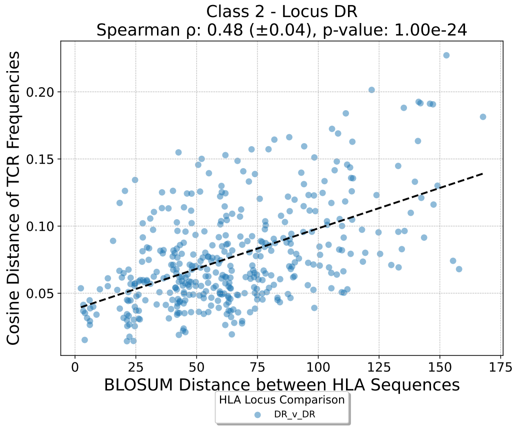


(f) Locus DR

Figure S7: Per locus positional analysis of the correlation between HLA sequence polymorphism and change in TCR V$\beta$ Preference.

#### C.3 Per Locus and Position

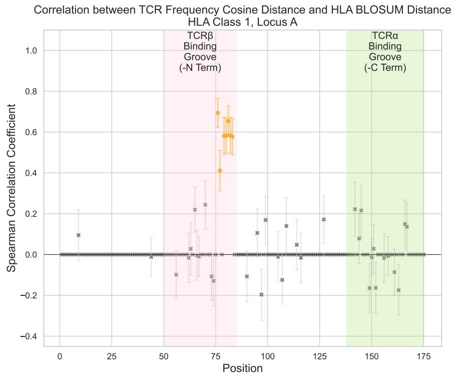


(a) HLA Locus A

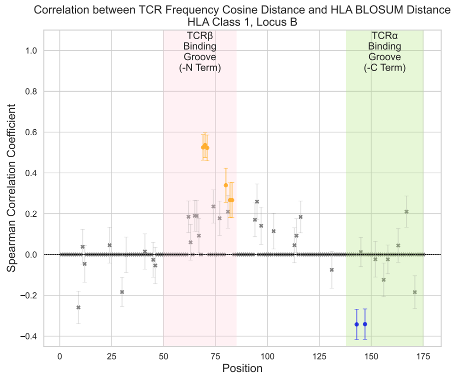


(b) HLA Locus B

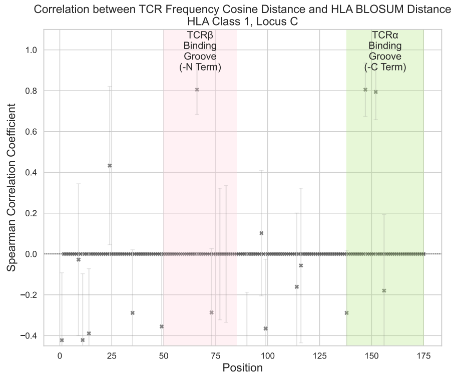


(c) HLA Locus C

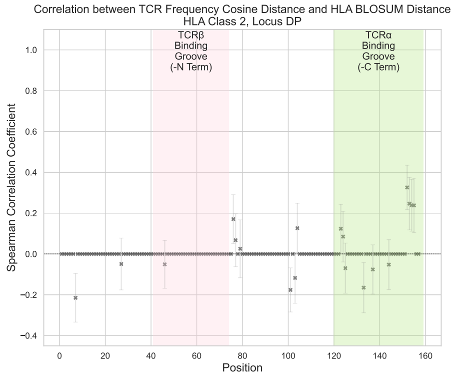


(d) HLA Locus DP

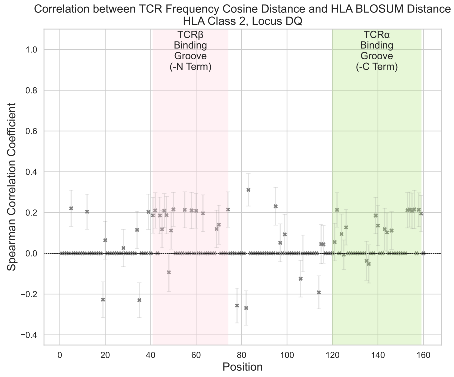


(e) HLA Locus DQ

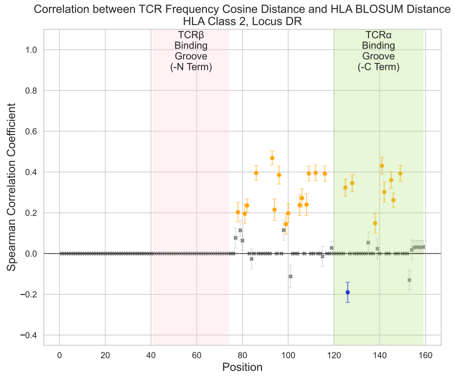


(f) HLA Locus DR

Figure S8: Positional Spearman $\rho$ of the correlation between change in HLA sequence as measured by BLOSUM distance and TCR V$\beta$ preference as measured by cosine distance, for each of the loci analyzed. We color in orange and blue the positions that pass the 0.01 significance threshold and False Discovery Rate (FDR) of 1%. Grey indicates no significance, orange indicates significant and positive correlation and blue indicates significant and negative correlation.

### D HLA Sequence Polymorphism and Peptide Motif JS Divergence correlation per Locus

Similar to Appendix C, here section we quantify the relationship between HLA polymorphism and changes in peptide motifs using Spearman correlations. We break this down into an overall correlation (Section D.1), per-locus correlation (Section C.2) and per-locus and position in the HLA sequence (Section D.3) to identify the main drivers of the patterns.

#### D.1 Overall

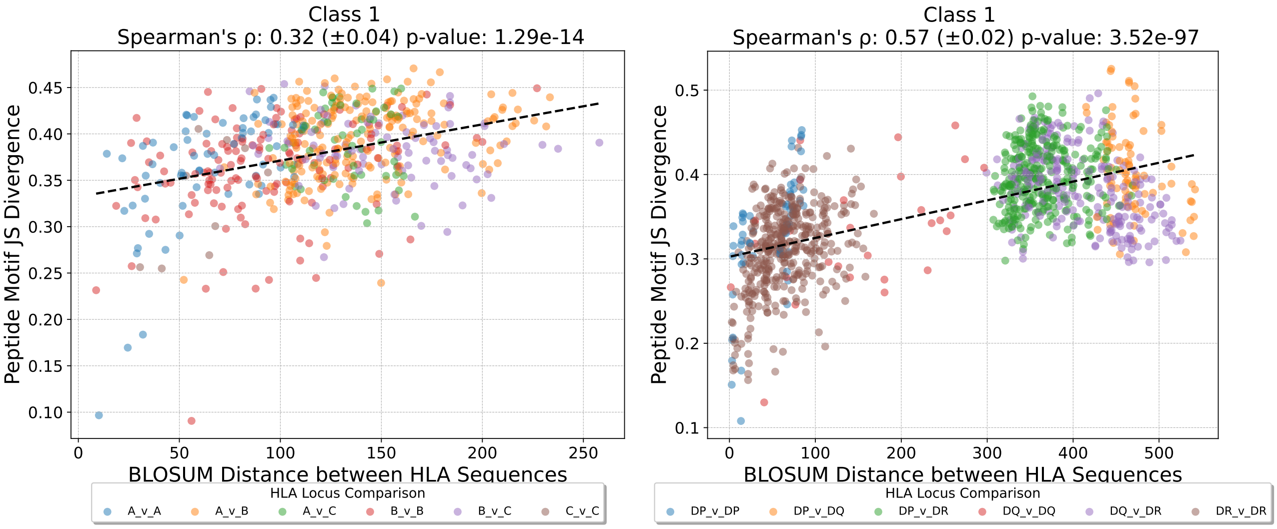


(a) HLA class I

(b) HLA class II

Figure S9: Relationship between HLA sequence polymorphism measured with BLOSUM distance and changes in Peptide Motifs measured with JS Divergence. The analysis is performed per HLA class and locus. See Table S3 for individual Spearman's $\rho$ values and Figure S10 for plots per locus.

| Class | Locus | Spearman $\rho$ (Bootstrap SD) | p-Value |
|---|---|---|---|
| 1 | all | 0.32 (0.04) | **1.29-14** |
| 2 | all | 0.57 (0.02) | **3.52-97** |
| 1 | A | 0.57 (0.09) | **4.31E-07** |
| 1 | B | 0.33 (0.08) | **9.76E-05** |
| 1 | C | 0.57 (0.28) | 8.97E-02 |
| 2 | DP | 0.80 (0.06) | **2.22E-13** |
| 2 | DQ | 0.48 (0.15) | **8.31E-03** |
| 2 | DR | 0.36 (0.05) | **5.25E-14** |

Table S3: Spearman $\rho$ coefficients and bootstrap standard deviation (SD) of HLA sequence polymorphism measured with BLOSUM distance and and changes in Peptide Motifs measured with JS Divergence for all loci and classes, using a significance threshold of 0.01. The p-Values of the tests that pass the threshold are in bold. See Figure S10 for a per locus plot.

#### D.2 Per Locus

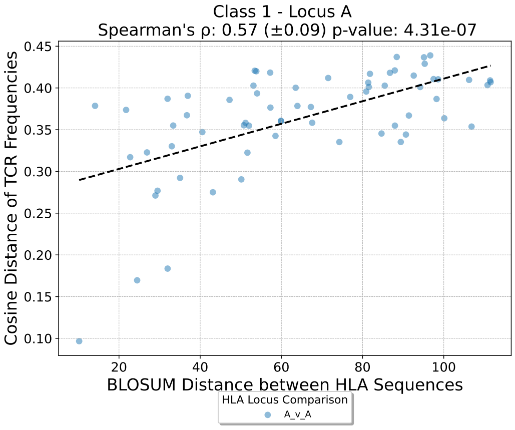


(a) Locus A

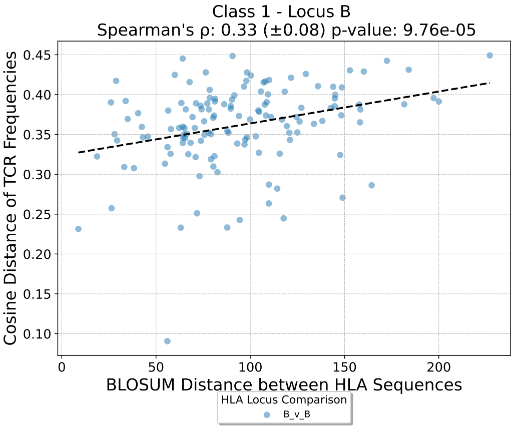


(b) Locus B

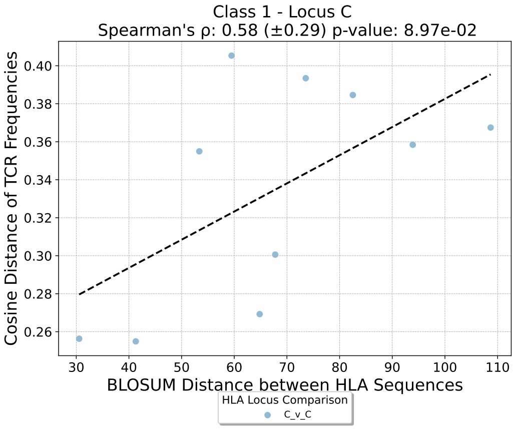


(c) Locus C

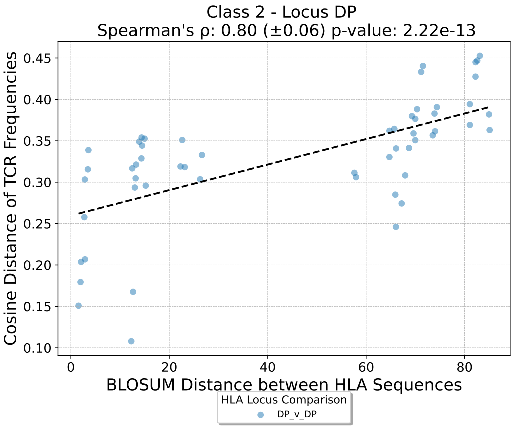


(d) Locus DP

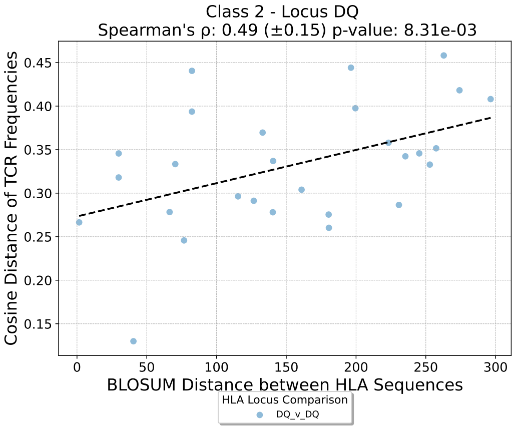


(e) Locus DQ

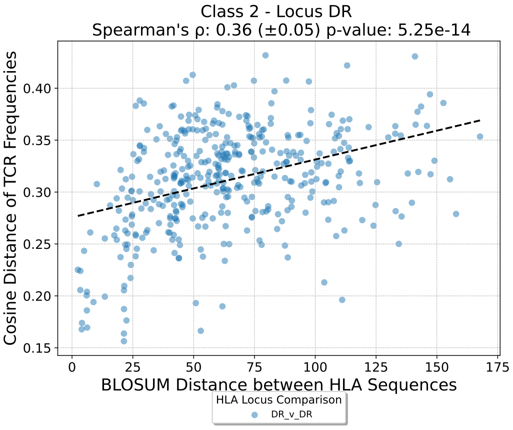


(f) Locus DR

Figure S10: Per locus positional analysis of the correlation between HLA sequence polymorphism and divergence in peptide motif

#### D.3 Per Locus and Per Position

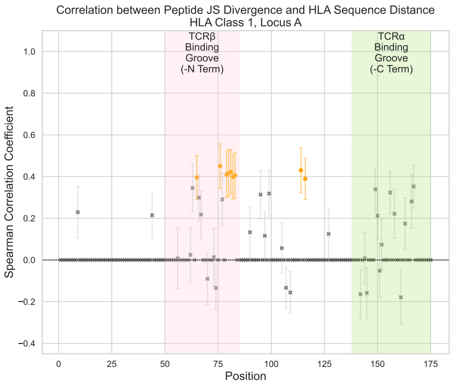


(a) Locus A

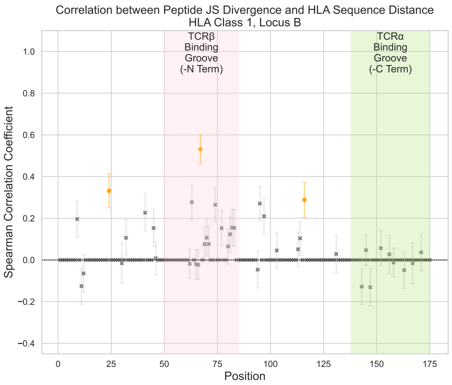


(b) Locus B

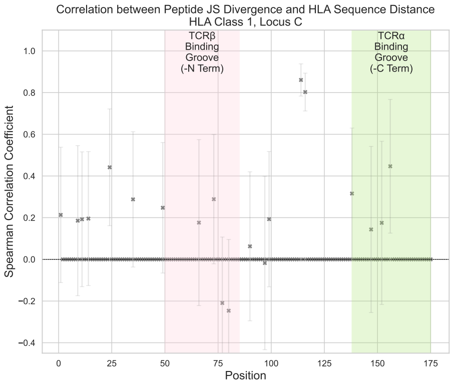


(c) Locus C

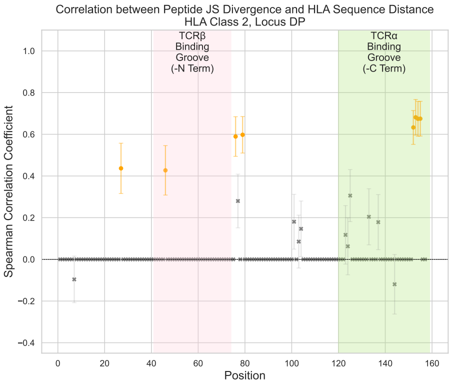


(d) Locus DP

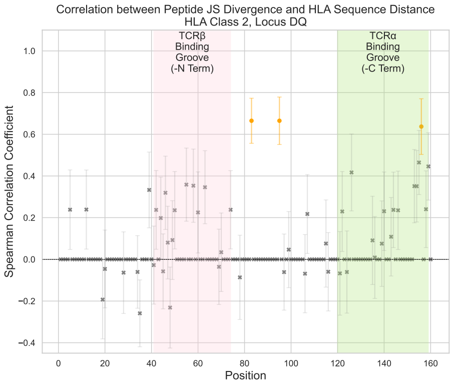


(e) Locus DQ

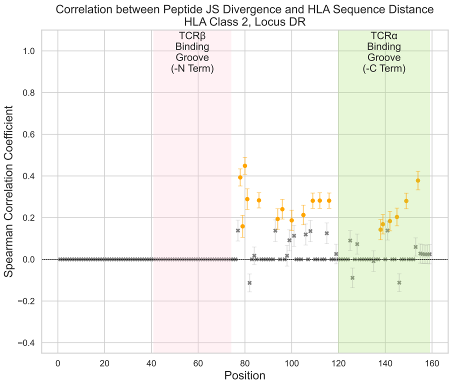


(f) Locus DR

Figure S11: Analysis of importance of HLA positions. Spearman $\rho$ of the correlation between change in HLA sequence as measured by BLOSUM distance and Peptide Motif as measured by JS Divergence, for each of the loci analysed. We color in orange and blue the positions that pass the 0.01 significance threshold and False Discovery Rate (FDR) of 1%. Grey indicates no significance, orange indicates significant and positive correlation and blue indicates significant and negative correlation.

### E TCR$\beta$ vs Peptide Disentanglement

To disentangle the contribution of individual HLA residues to each signal, in Figure S12, we plot the correlation coefficient for each position with TCR$\beta$ divergence (x-axis) and peptide divergence (y-axis). HLA positions above the identity line (red) are more strongly correlated with peptide differences, while those below are more associated with TCR$\beta$ divergence.

In Section E.2, we list the exact correlation coefficients and standard deviation of each of the plotted position, while in Section E.3, we visualize these positions onto the 3D structures of the HLAs.

#### E.1 Disentangling significant positions by importance for TCR$\beta$ and Peptide

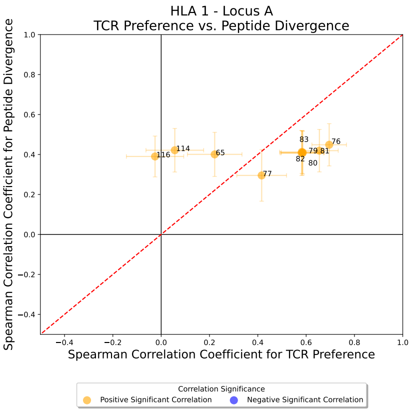


(a) Locus A

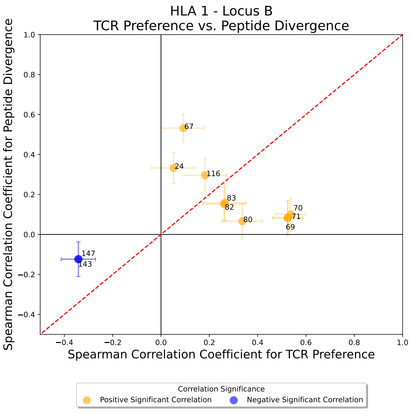


(b) Locus B

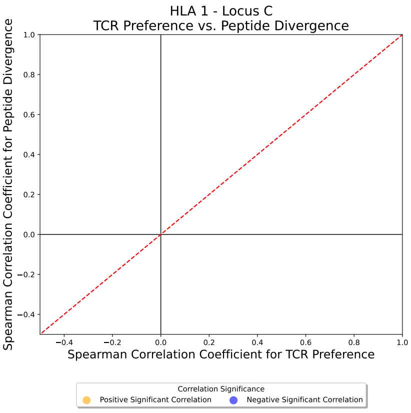


(c) Locus C

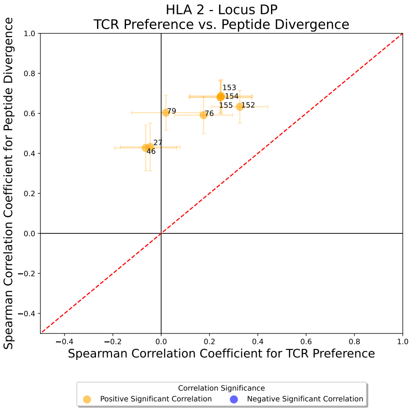


(d) Locus DP

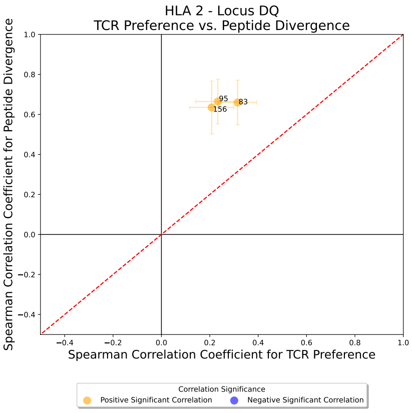


(e) Locus DQ

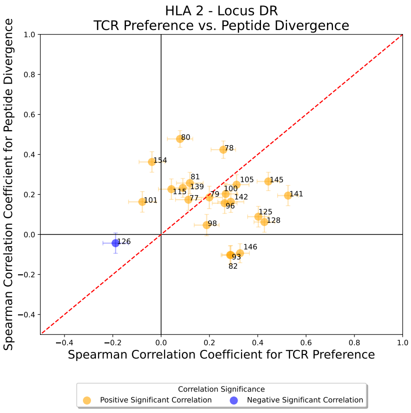


(f) Locus DR

Figure S12: Visualization Spearman $\rho$ of significant positions in the HLA sequence towards the TCR$\beta$ or the peptide. The red line indicates equal strength of correlation. Blue indicates negative correlation, while orange is positive correlation.

#### E.2 Positions Per HLA Locus

| Position | Bootstrap Corr (TCR$\beta$) | Bootstrap Std Dev (TCR$\beta$) | p-value (TCR$\beta$) | Bootstrap Corr (peptide) | Bootstrap Std Dev (peptide) | p-value (peptide) |
|---|---|---|---|---|---|---|
| 65 | 0.225 | 0.111 | 0.306 | 0.396 | 0.106 | 0.005 |
| 76 | 0.695 | 0.071 | 0.0 | 0.445 | 0.107 | 0.005 |
| 77 | 0.412 | 0.103 | 0.003 | 0.29 | 0.125 | 0.044 |
| 79 | 0.582 | 0.086 | 0.0 | 0.403 | 0.107 | 0.005 |
| 80 | 0.582 | 0.089 | 0.0 | 0.403 | 0.103 | 0.005 |
| 81 | 0.652 | 0.081 | 0.0 | 0.414 | 0.104 | 0.005 |
| 82 | 0.582 | 0.089 | 0.0 | 0.403 | 0.112 | 0.005 |
| 83 | 0.582 | 0.085 | 0.0 | 0.403 | 0.105 | 0.005 |
| 114 | 0.05 | 0.12 | 0.941 | 0.421 | 0.103 | 0.005 |
| 116 | -0.015 | 0.124 | 0.98 | 0.389 | 0.101 | 0.006 |

Table S4: Locus A

| Position | Bootstrap Corr (TCR$\beta$) | Bootstrap Std Dev (TCR$\beta$) | p-value (TCR$\beta$) | Bootstrap Corr (peptide) | Bootstrap Std Dev (peptide) | p-value (peptide) |
|---|---|---|---|---|---|---|
| 24 | 0.049 | 0.093 | 0.737 | 0.333 | 0.078 | 0.001 |
| 67 | 0.094 | 0.086 | 0.441 | 0.532 | 0.073 | 0.0 |
| 69 | 0.525 | 0.06 | 0.0 | 0.078 | 0.084 | 0.664 |
| 70 | 0.538 | 0.062 | 0.0 | 0.098 | 0.087 | 0.51 |
| 71 | 0.525 | 0.062 | 0.0 | 0.078 | 0.084 | 0.664 |
| 80 | 0.338 | 0.083 | 0.0 | 0.06 | 0.083 | 0.803 |
| 82 | 0.265 | 0.084 | 0.009 | 0.156 | 0.086 | 0.234 |
| 83 | 0.265 | 0.088 | 0.009 | 0.156 | 0.084 | 0.234 |
| 116 | 0.178 | 0.083 | 0.079 | 0.296 | 0.082 | 0.006 |
| 143 | -0.343 | 0.071 | 0.0 | -0.126 | 0.087 | 0.358 |
| 147 | -0.343 | 0.073 | 0.0 | -0.126 | 0.088 | 0.358 |

Table S5: Locus B

| Position | Bootstrap Corr (TCR$\beta$) | Bootstrap Std Dev (TCR$\beta$) | p-value (TCR$\beta$) | Bootstrap Corr (peptide) | Bootstrap Std Dev (peptide) | p-value (peptide) |
|---|---|---|---|---|---|---|
| 27 | -0.05 | 0.123 | 0.727 | 0.426 | 0.12 | 0.003 |
| 46 | -0.05 | 0.124 | 0.727 | 0.426 | 0.113 | 0.003 |
| 76 | 0.169 | 0.122 | 0.448 | 0.587 | 0.093 | 0.0 |
| 79 | 0.031 | 0.135 | 0.805 | 0.596 | 0.088 | 0.0 |
| 152 | 0.319 | 0.113 | 0.17 | 0.633 | 0.082 | 0.0 |
| 153 | 0.241 | 0.13 | 0.244 | 0.679 | 0.082 | 0.0 |
| 154 | 0.241 | 0.13 | 0.244 | 0.679 | 0.084 | 0.0 |
| 155 | 0.241 | 0.13 | 0.244 | 0.679 | 0.082 | 0.0 |

Table S6: Locus DP

| Position | Bootstrap Corr (TCR$\beta$) | Bootstrap Std Dev (TCR$\beta$) | p-value (TCR$\beta$) | Bootstrap Corr (peptide) | Bootstrap Std Dev (peptide) | p-value (peptide) |
|---|---|---|---|---|---|---|
| 83 | 0.315 | 0.08 | 0.024 | 0.659 | 0.114 | 0.003 |
| 95 | 0.228 | 0.09 | 0.056 | 0.659 | 0.113 | 0.003 |
| 156 | 0.209 | 0.091 | 0.056 | 0.633 | 0.13 | 0.005 |

Table S7: Locus DQ

| Position | Bootstrap Corr (TCR$\beta$) | Bootstrap Std Dev (TCR$\beta$) | p-value (TCR$\beta$) | Bootstrap Corr (peptide) | Bootstrap Std Dev (peptide) | p-value (peptide) |
|---|---|---|---|---|---|---|
| 78 | 0.204 | 0.048 | 0.0 | 0.391 | 0.043 | 0.0 |
| 79 | 0.111 | 0.053 | 0.037 | 0.159 | 0.052 | 0.003 |
| 80 | 0.066 | 0.048 | 0.256 | 0.447 | 0.04 | 0.0 |
| 81 | 0.197 | 0.049 | 0.0 | 0.289 | 0.047 | 0.0 |
| 82 | 0.237 | 0.032 | 0.0 | -0.112 | 0.044 | 0.037 |
| 86 | 0.396 | 0.035 | 0.0 | 0.283 | 0.038 | 0.0 |
| 93 | 0.47 | 0.034 | 0.0 | 0.135 | 0.05 | 0.012 |
| 94 | 0.214 | 0.053 | 0.0 | 0.193 | 0.05 | 0.0 |
| 96 | 0.387 | 0.043 | 0.0 | 0.243 | 0.049 | 0.0 |
| 99 | 0.147 | 0.048 | 0.005 | 0.092 | 0.048 | 0.095 |
| 100 | 0.195 | 0.045 | 0.0 | 0.188 | 0.049 | 0.0 |
| 105 | 0.24 | 0.051 | 0.0 | 0.213 | 0.047 | 0.0 |
| 106 | 0.271 | 0.047 | 0.0 | 0.121 | 0.048 | 0.025 |
| 108 | 0.238 | 0.052 | 0.0 | 0.136 | 0.05 | 0.012 |
| 109 | 0.396 | 0.037 | 0.0 | 0.283 | 0.04 | 0.0 |
| 112 | 0.396 | 0.036 | 0.0 | 0.283 | 0.037 | 0.0 |
| 116 | 0.396 | 0.037 | 0.0 | 0.283 | 0.038 | 0.0 |
| 125 | 0.324 | 0.045 | 0.0 | 0.088 | 0.048 | 0.107 |
| 126 | -0.19 | 0.051 | 0.0 | -0.088 | 0.045 | 0.107 |
| 128 | 0.348 | 0.04 | 0.0 | 0.072 | 0.046 | 0.198 |
| 138 | 0.145 | 0.049 | 0.006 | 0.144 | 0.049 | 0.008 |
| 139 | 0.024 | 0.051 | 0.663 | 0.167 | 0.046 | 0.002 |
| 141 | 0.432 | 0.039 | 0.0 | 0.137 | 0.048 | 0.012 |
| 142 | 0.306 | 0.047 | 0.0 | 0.184 | 0.049 | 0.001 |
| 145 | 0.363 | 0.04 | 0.0 | 0.204 | 0.045 | 0.0 |
| 146 | 0.265 | 0.036 | 0.0 | -0.109 | 0.043 | 0.043 |
| 149 | 0.396 | 0.038 | 0.0 | 0.283 | 0.039 | 0.0 |
| 154 | 0.023 | 0.048 | 0.663 | 0.378 | 0.046 | 0.0 |

Table S8: Locus DR

#### E.3 Visualizing Disentangled Positions on the 3D structure

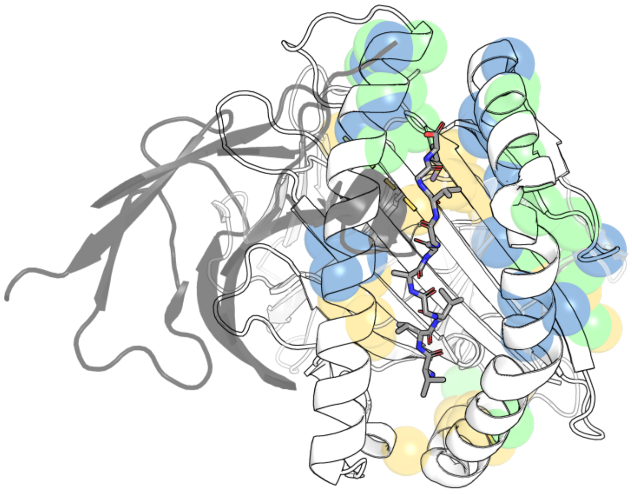

(a) All Loci

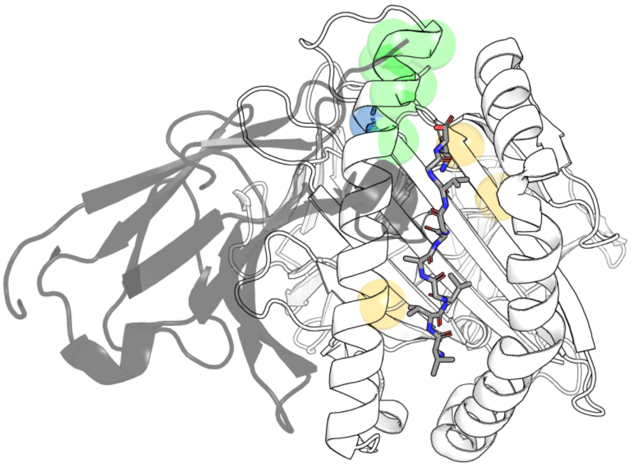

(b) Locus A

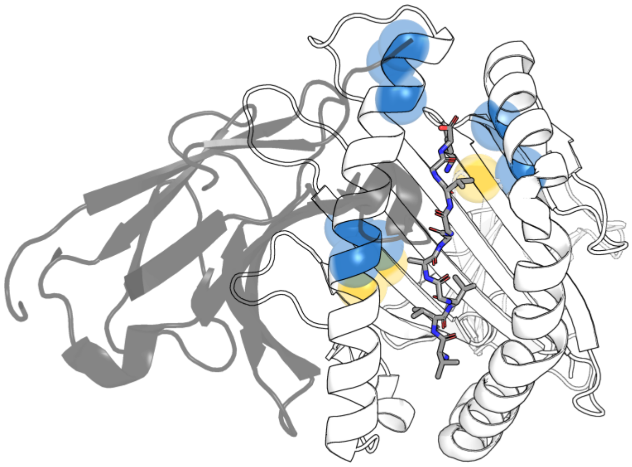

(c) Locus B

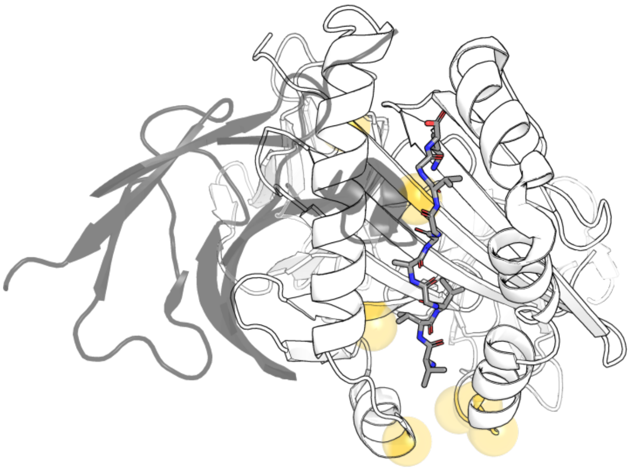

(d) Locus DP

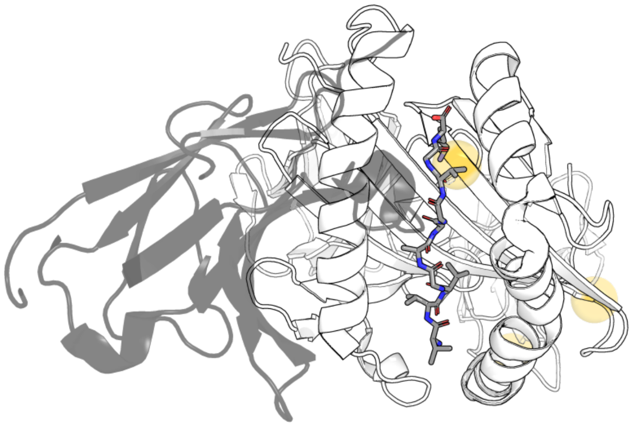

(e) Locus DQ

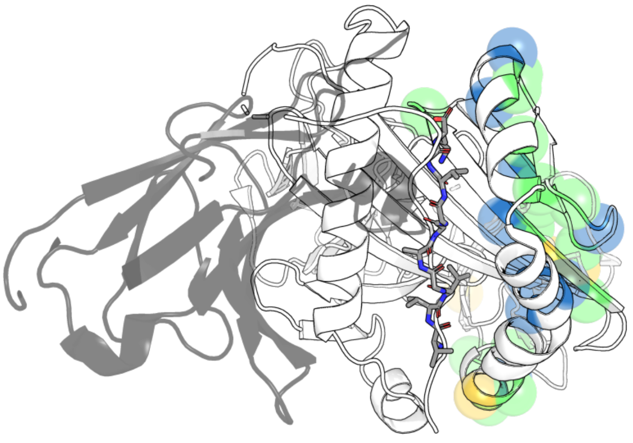

(f) Locus DR

Figure S13: 3D Plot of significant HLA positions colored by importance for the peptide (yellow), TCR$\beta$ (blue) or both (green). Figure A visualizes all the positions relevant for all loci on an HLA-A structure (PDB: 7OW5). In grey the TCR$\beta$ contacting the $\alpha_1$ helix.

#### F Visualising TCR$\beta$ Y46 Y48 and E54 contacts with HLA

Some of the significant HLA positions identified (65 and 68 in the $\alpha$1 helix) are within close contact[1] of a conserved CDR2 motif (Y46, Y48 and E54), shared across species (Scott-Browne et al., 2011). In Figure Figure S14, we overlap the disentangled positions from Figure S13 and Positions 65 and 68 in orange. In our analysis, position 68 was significant for the peptide and both positions 65 and 68 are near positions significant for both the TCR and peptide.

Mutational studies targeting this TCR motif have shown impaired thymic selection and decreased development of the TCR repertoire, supporting a model where germline-encoded TCR-HLA contacts may guide thymic selection, ultimately shaping the mature repertoire toward effective pHLA recognition (Scott-Browne et al., 2011; Krovi et al., 2019; Sharon et al., 2016).

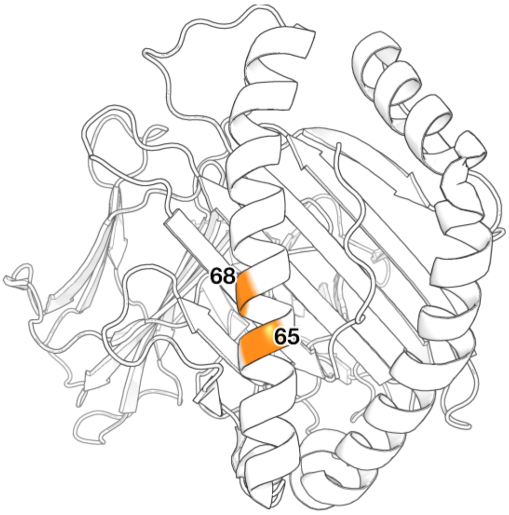


(a) HLA class I conserved contact positions.

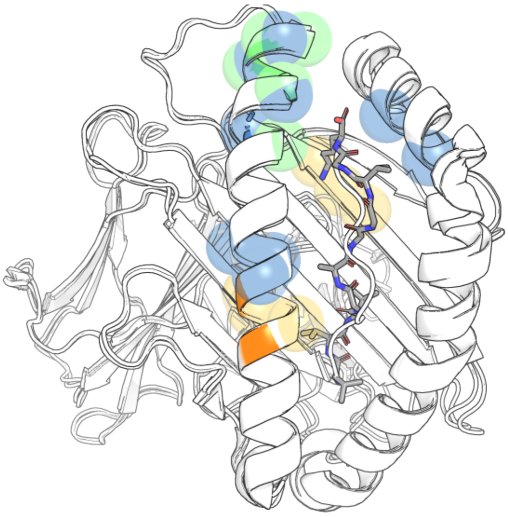

(b) HLA class I significant positions (from Figure S13) and conserved contact positions (orange).

Figure S14: 3D Plot of significant HLA positions colored by importance for the peptide (yellow), TCR$\beta$ (blue) or both (green), visualised on an HLA-A structure (PDB: 7OW5). In orange, are position 65 and 68 of the HLA, in contact with the conserved CDR2 motif Y46 Y48 and E54 (PDB: 1LP9).

[1]Under 4 Å

### G Peptide Motif JS Divergence and TCR V$\beta$ Preference correlation

To assess whether similarity in peptide motifs contributes to observed V$\beta$ gene usage patterns across HLAs, we computed pairwise correlations between peptide motif divergence and TCR V$\beta$ preference. For each HLA pair, peptide motifs were compared using JS divergence between PWMs, while V$\beta$ gene usage differences with cosine distance of normalize.

In Section G.1, we assess overall and class-specific correlations across all HLAs. In Section G.2, we break down the analysis by locus to identify which HLA loci are more strongly correlated. Finally, in Section G.3, we examine per-peptide position contributions to the correlations.

#### G.1 Overall

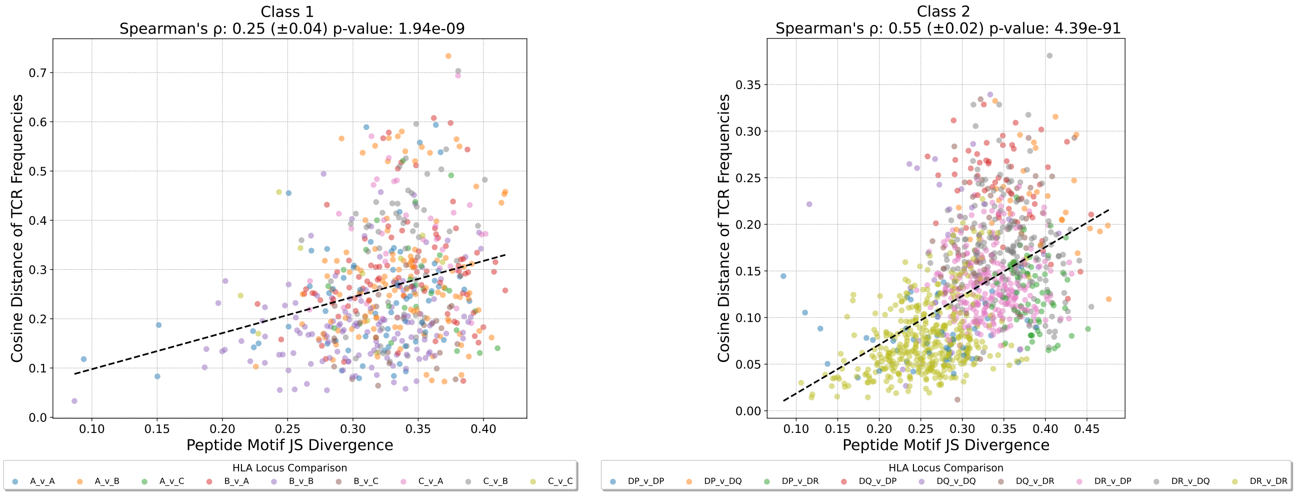


(a) HLA class I

(b) HLA class II

Figure S15: Relationship between changes in TCR V$\beta$ preference measured with cosine distance and changes in Peptide Motifs measured with JS Divergence. Analysis is performed per class and per locus. See Table S9 for Spearman's $\rho$ values nd Figure S7 for plots per locus.

| Class | Locus | Spearman $\rho$ (Bootstrap SD) | p-Value |
|---|---|---|---|
| 1 | all | 0.25 (0.04) | **1.08E-09** |
| 2 | all | 0.52 (0.02) | **1.04E-74** |
| 1 | A | 0.25 (0.12) | 3.61E-02 |
| 1 | B | 0.23 (0.08) | **7.25E-03** |
| 1 | C | 0.19 (0.33) | 6.27E-01 |
| 2 | DP | 0.07 (0.15) | 6.02E-01 |
| 2 | DQ | 0.04 (0.19) | 8.38E-01 |
| 2 | DR | 0.35 (0.05) | **9.44E-13** |

Table S9: Spearman $\rho$ coefficients and Standard Deviation (SD) of changes in TCR V$\beta$ preference as measured by cosine distance and changes in Peptide Motifs as measured by JS Divergence for all loci and classes, with significance threshold of 0.01. The p-Values of the tests that pass the threshold are in bold. For a per locus plot refer to Figure S16.

#### G.2 Per Locus

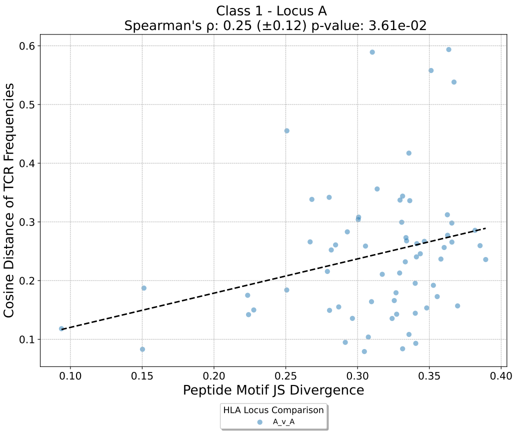


(a) Locus A

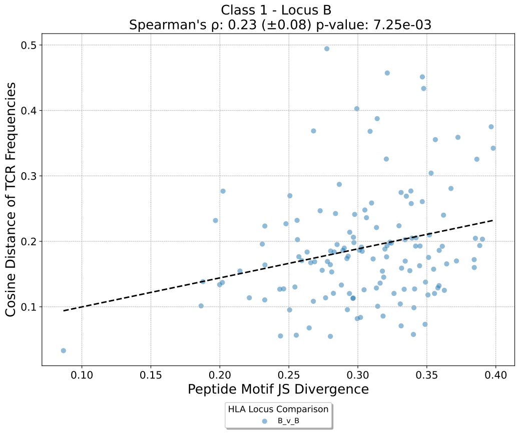


(b) Locus B

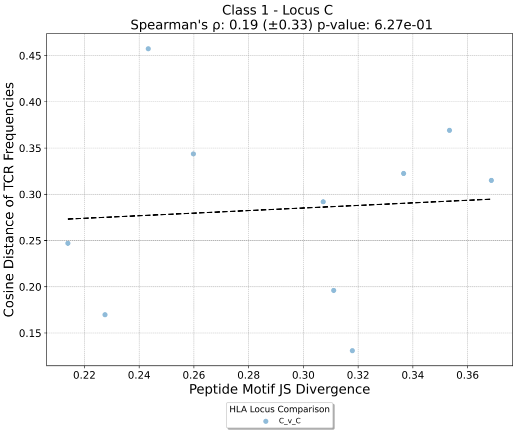


(c) Locus C

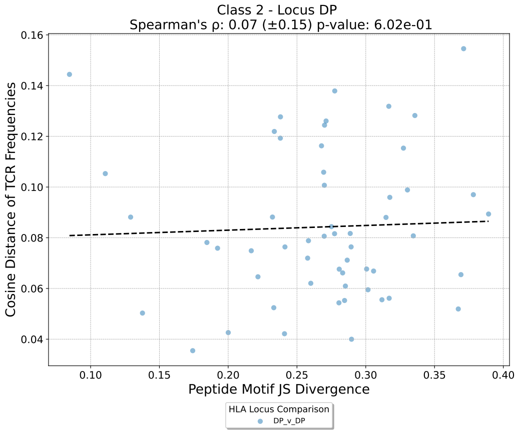


(d) Locus DP

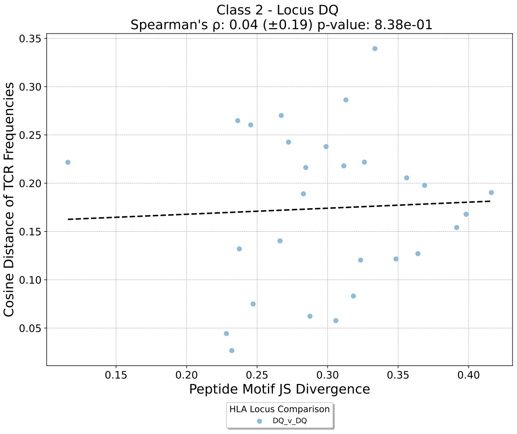


(e) Locus DQ

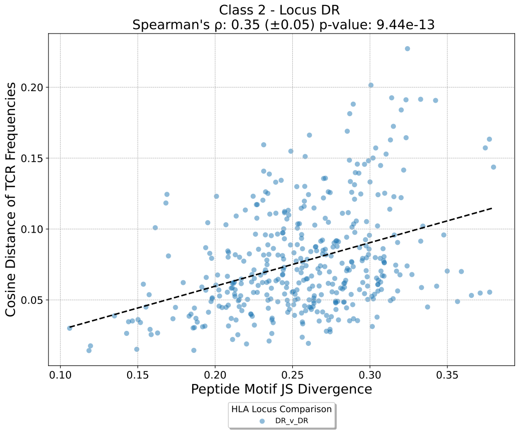


(f) Locus DR

Figure S16: Per locus positional analysis of the correlation between divergence in peptide motif and change in TCR V$\beta$ Preference.

#### G.3 Per Locus and per position

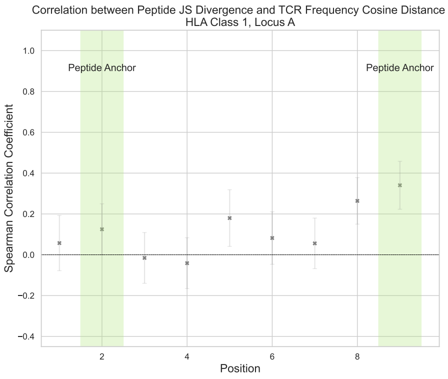


(a) Locus A

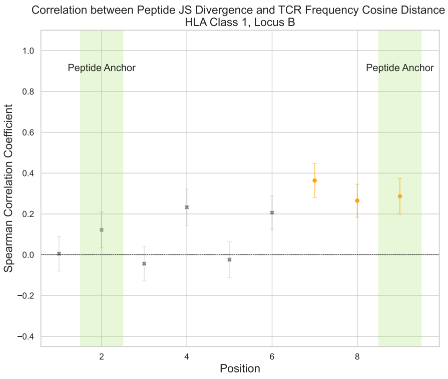


(b) Locus B

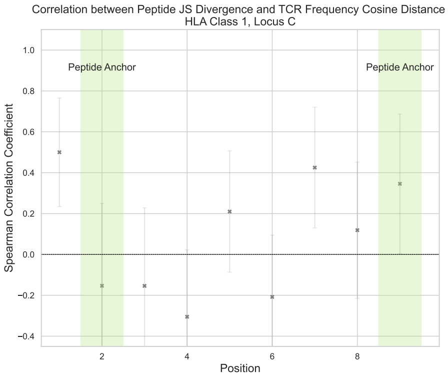


(c) Locus C

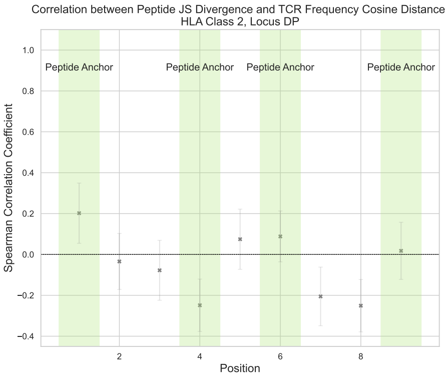


(d) Locus DP

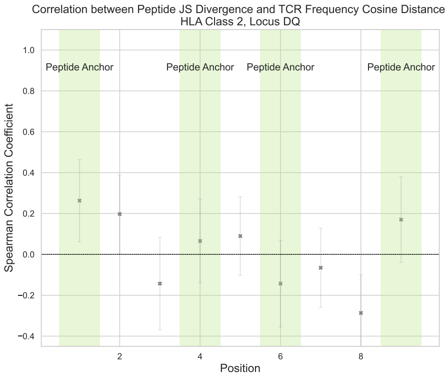


(e) Locus DQ

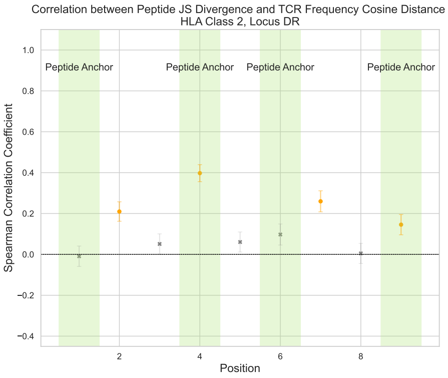


(f) Locus DR

Figure S17: Analysis of importance of Peptide positions in each HLA Locus. Spearman $\rho$ of the correlation between change in Peptide Motif measured with JS divergence TCR V$\beta$ preference as measured by cosine distances, for each of the loci analysed. We color in orange the positions that pass the 0.01 significance threshold and False Discovery Rate (FDR) of 1%. Grey indicates no significance, orange indicates significant correlation.

#### H Quantitative Analysis of HLA Restriction Breadth

To validate the distinction between "specialist" (restricted) and "generalist" (promiscuous) V$\beta$ usage, we quantified the breadth of HLA restriction for each locus using normalized Shannon entropy ($H_{\text{norm}} = H/\ln N$). This metric scales from 0 (exclusive restriction to a single allele) to 1 (uniform interaction across all alleles). Figure S18 summarizes the restriction breadth for each locus.

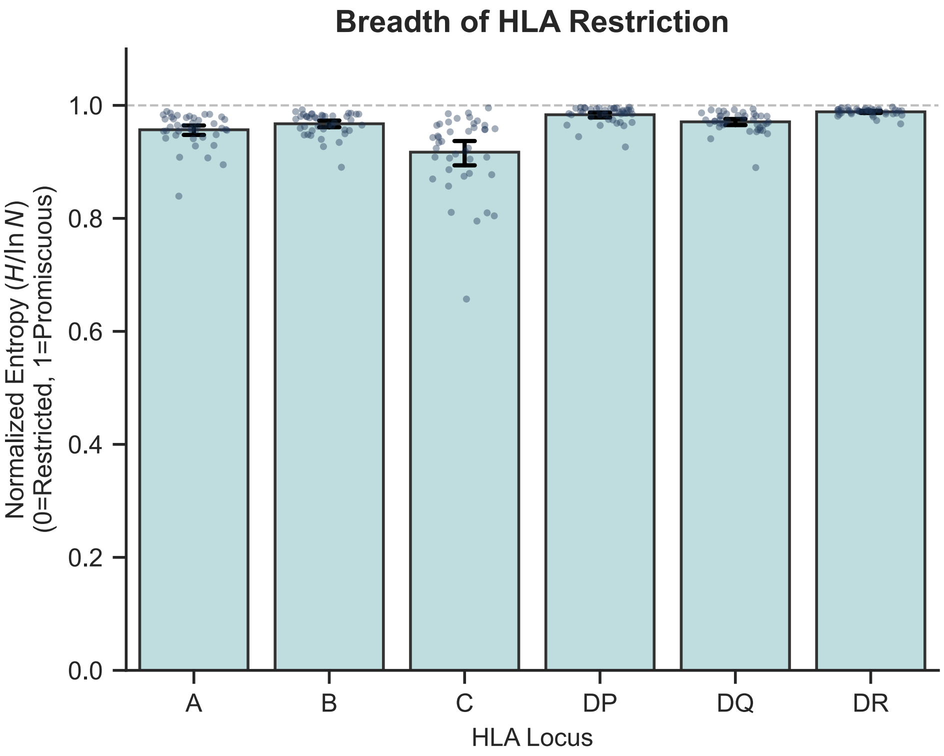


Figure S18: Comparison of V$\beta$ restriction breadth across HLA loci. Bars represent the mean normalized entropy ($H/\ln N$) for each locus, with error bars indicating the standard deviation. Individual points represent specific V$\beta$ genes. A value of 1.0 (dashed line) indicates uniform promiscuity (generalist), while lower values indicate increasing HLA restriction (specialist). HLA-C shows the lowest mean and highest variance, indicative of a subset of highly restricted interactions.

To confirm these differences were statistically significant, we performed pairwise Mann-Whitney U tests between the normalized entropy distributions of all loci, applying a Bonferroni correction for multiple comparisons (Table S10).

| Locus 1 | Locus 2 | MW Stat | P-Value | Adj. P-Value | Sig. (< 0.05) | Sig. (< 0.001) |
|---|---|---|---|---|---|---|
| C | DR | 66 | 7.07E-13 | 1.06E-11 | TRUE | TRUE |
| A | DR | 72 | 1.06E-12 | 1.59E-11 | TRUE | TRUE |
| C | DP | 143 | 1.02E-10 | 1.53E-09 | TRUE | TRUE |
| B | DR | 145 | 1.15E-10 | 1.73E-09 | TRUE | TRUE |
| DQ | DR | 221 | 9.43E-09 | 1.42E-07 | TRUE | TRUE |
| A | DP | 229 | 1.46E-08 | 2.19E-07 | TRUE | TRUE |
| C | DQ | 300 | 5.50E-07 | 8.25E-06 | TRUE | TRUE |
| B | DP | 339 | 3.38E-06 | 5.07E-05 | TRUE | TRUE |
| B | C | 1325 | 7.17E-06 | 0.00 | TRUE | TRUE |
| DP | DQ | 1276 | 5.48E-05 | 0.00 | TRUE | TRUE |
| A | C | 1185 | 0.00 | 0.02 | TRUE | FALSE |
| A | DQ | 562 | 0.01 | 0.15 | FALSE | FALSE |
| A | B | 621 | 0.04 | 0.63 | FALSE | FALSE |
| B | DQ | 754 | 0.43 | 1.00 | FALSE | FALSE |
| DP | DR | 749 | 0.40 | 1.00 | FALSE | FALSE |

Table S10: Pairwise Mann-Whitney U test results comparing the normalized entropy distributions of V$\beta$ restriction across HLA loci. P-values were adjusted for multiple comparisons using the Bonferroni correction.
